# AT1R-Selective Negative Allosteric Modulator Targeting the Epitope for Pathogenic Autoantibody Activation

**DOI:** 10.64898/2026.10.12.758523

**Authors:** Khuraijam Dhanachandra Singh, Triveni R. Pardhi, Terri J. Harford, Luxmichan Laishram, Tarun Ravi, Russell Desnoyer, Anjli P. Maroo, W. H. Wilson Tang, Sadashiva S. Karnik

**Affiliations:** Heart, Blood and Kidney Research Department; Fairview Hospital; Cardiovascular Medicine Department, Cleveland Clinic Lerner College of Medicine at Case Western Reserve University, Cleveland, OH 44195, U.S.A

**Keywords:** AT1R, AngII, AT1R-AAb, NAM, Allosteric ligand, GPCR, blood pressure, pharmacokinetics, antihypertensives

## Abstract

A major therapeutic strategy for managing cardiovascular and renal diseases is the use of angiotensin receptor blockers (ARBs). These drugs block both pathogenic and physiological signaling by the angiotensin II type 1 receptor (AT1R) by preventing angiotensin II (AngII) binding to orthosteric pocket in transmembrane portion of AT1R. However, in many autoimmune diseases, AT1R activation is chronically driven by pathogenic autoantibody binding to a distinct extracellular surface located allosteric site. Currently, there are no allosteric inhibitors of AT1R available to mechanistically prevent the autoantibodies and preserve the orthosteric hormonal functions of AngII. Here, we report the structure-guided hit-to-lead optimization of TP1, a first-in-class AT1R allosteric chemotype and negative allosteric modulator (NAM) with improved potency and drug-like properties. Computational docking and site-directed mutagenesis identified a TP1 binding mode that is distinct from the orthosteric-site binding of ARBs and AngII. Synthetically pure TP1 significantly reduced the capacity of AT1R to bind AngII without affecting AngII binding to AT2R, demonstrating an AT1R selective effect. TP1 reduced AngII-induced calcium signaling and migration in AT1R-expressing cells, arterial contraction, and increases in blood pressure in AngII-infused mice, demonstrating its activity as a NAM. Crucially, TP1 blocked binding of pathogenic autoantibodies to AT1R-expressing cells from the serum of patients with preeclampsia and dilated cardiomyopathy, a function distinct from ARBs. Together, these findings establish TP1 as a proof-of-concept, AT1R-selective NAM chemotype with therapeutic potential for AT1R autoantibody-driven conditions such as preeclampsia, where ARBs are strictly contraindicated. Developing AT1R-targeted NAM antihypertensive drugs may offer therapeutic advantages over ARBs and expand the pharmacological toolkit for this important CVD receptor.

## Introduction

The angiotensin II type 1 receptor (AT1R) is a G protein-coupled receptor (GPCR) and one of the most important drug targets in cardiovascular medicine. AT1R mediates the hormonal actions of angiotensin II (AngII), an endogenous octapeptide in the renin– angiotensin system that plays essential roles in heart and blood vessel contraction and in regulating fluid and electrolyte balance in the kidneys. However, excessive or dysregulated AT1R signaling also contributes to inflammation, fibrosis, autoimmunity, and metabolic disorders^1^.

Angiotensin receptor blockers (ARBs) are currently a cornerstone of antihypertensive, reno-protective, cardioprotective, antifibrotic, and anti-inflammatory therapies. ARBs act as classical antagonists or inverse agonists by preventing AngII from binding to the orthosteric pocket of AT1R^1–6^. For decades, drug development has focused on the same orthosteric pocket of AT1R including the efforts to develop β-arrestin-biased AT1R orthosteric agonists, which displace AngII while selectively modifying downstream signaling^7^. As a result, the field has relied on a orthosteric site-restricted pharmacological mechanism, limiting the potential for greater therapeutic versatility^8–10^. ARB treatment, which blocks the physiological effects of AngII, may have important clinical drawbacks, including reduced contraction of the heart and blood vessels and impaired innate ability of blood vessels to repair and regenerate. Furthermore, preservation of physiological AngII signaling is essential in developmental settings, including fetal kidney and liver development. Therefore, complete pharmacological blockade of AT1R during pregnancy can be harmful, as demonstrated by the well-documented increase of fetopathy associated with ARB exposure^11,12^.

The allosteric ligands, in contrast, bind to topographically distinct receptor sites and fine-tune the activity, offering an innovative approach for drug development^13–15^. Unlike orthosteric ligands, allosteric ligands do not directly compete with endogenous hormones, allowing endogenic signaling to be preserved in a spatially and temporally regulated manner. They also offer greater receptor selectivity, improved safety, and probe-independent pharmacological effects^13–16^. These unique properties make allosteric modulators attractive therapeutic candidates. Although allosteric drugs have successfully advanced to the clinic for several GPCR families^15^, allosteric modulation of AT1R has remained largely unexplored. Addressing this knowledge gap, we previously identified a druggable allosteric pocket in the extracellular loop 2 (ECL2) of AT1R and reported initial hit compounds (DCP1-18) that act as allosteric modulators of response of AT1R to concentrations of AngII^17–19^. The DCP-hit compounds do not directly compete with AngII, reduce excessive AT1R activation and alter the receptor conformation in a way that prevents surrogate autoantibodies from binding to and activating the AT1R. However, the poor efficacy and limited drug-like properties of DCP-compounds restrict their utility for therapeutic development.

In diseases like preeclampsia (PE)^20,21^, Systemic Sclerosis^22,23^, Dilated Cardiomyopathy (DCM)^24,25^, Organ Transplant Rejection^26–29^, Lupus Nephritis^30,31^, Malignant Hypertension^27,32,33^, Postural Tachycardia Syndrome (POTS)^34,35^, and Primary Aldosteronism^36^, the patients produce autoantibodies against ECL2 of AT1R (AT1R-AAbs). Unlike normal antibodies, the AT1R-AAbs can activate the receptor continuously, leading to excessive signaling that underlies disease pathology. We discovered that the ECL2 allosteric pocket for DCP-compounds overlaps the epitope recognized by AT1R-AAbs^17,26,37,38^. Thus, targeting this allosteric site could be a promising way to develop drugs that block AT1R activation by AT1R-AAb in many autoimmune diseases.

As a proof of concept, synthetic ECL2-epitope peptides were shown to block AT1R-AAb-mediated PE pathology in preclinical models^39,40^. However, their clinical translation is hindered by the pharmacokinetic instability typical of peptide biologics and the well-established risk of boosting the antibody titer in patients, underscoring the need for the development of more potent small-molecule alternatives^41^.

Here, we report on a novel lead chemotype, TP1, designed to bind to the allosteric site in ECL2 of AT1R. TP1 was developed by optimizing the pharmacophore features of DCP1 hit compounds through structure-guided drug design. It is based on a new amino-pyridino-thiazole structure and acts as a negative allosteric modulator (NAM) of AT1R. Using structural, pharmacological, physiological, and pharmacokinetic studies, we identified how TP1 alters AT1R activity and demonstrated its efficacy across in vitro and in vivo models. Importantly, we found that TP1 blocked AT1R-AAb from patients with preeclampsia (PE) and dilated cardiomyopathy (DCM). This suggests that TP1 may target a common mechanism involved in AT1R-AAb-related diseases. Overall, these efforts advance AT1R allosteric drug development and identify TP1 as a promising new drug candidate.

## Results

### Structure-Guided Hit-to-Lead Optimization Yields TP1

Our previous proof-of-concept studies identified DCP1-3 and DCP1-16 as small-molecule NAMs of AT1R with low efficacy^17^. For example, these compounds reduced saturation binding (B_max_) of the partial agonist AngIV but did not affect the binding of full agonist AngII. Their NAM activity at AT1R signaling was observed in the 1-100 µM concentration range. Both compounds share a common architecture centered on a trisubstituted cyclopentane core bearing a tetrahydropyranyl amide cap, a methylated ethanolamine tail, a free hydroxyl group, and a halogenated aryl ether arm, monofluorophenyl in DCP1-3 and difluorophenyl in DCP1-16, with DCP1-16 additionally bearing a methoxyphenyl substituent on the amine tail (**Fig. 1**). Thus, low efficacy, structural complexity, multiple stereocenters, and halogenated aryl substituents made these compounds poor candidates for further development.

**Figure 1.**
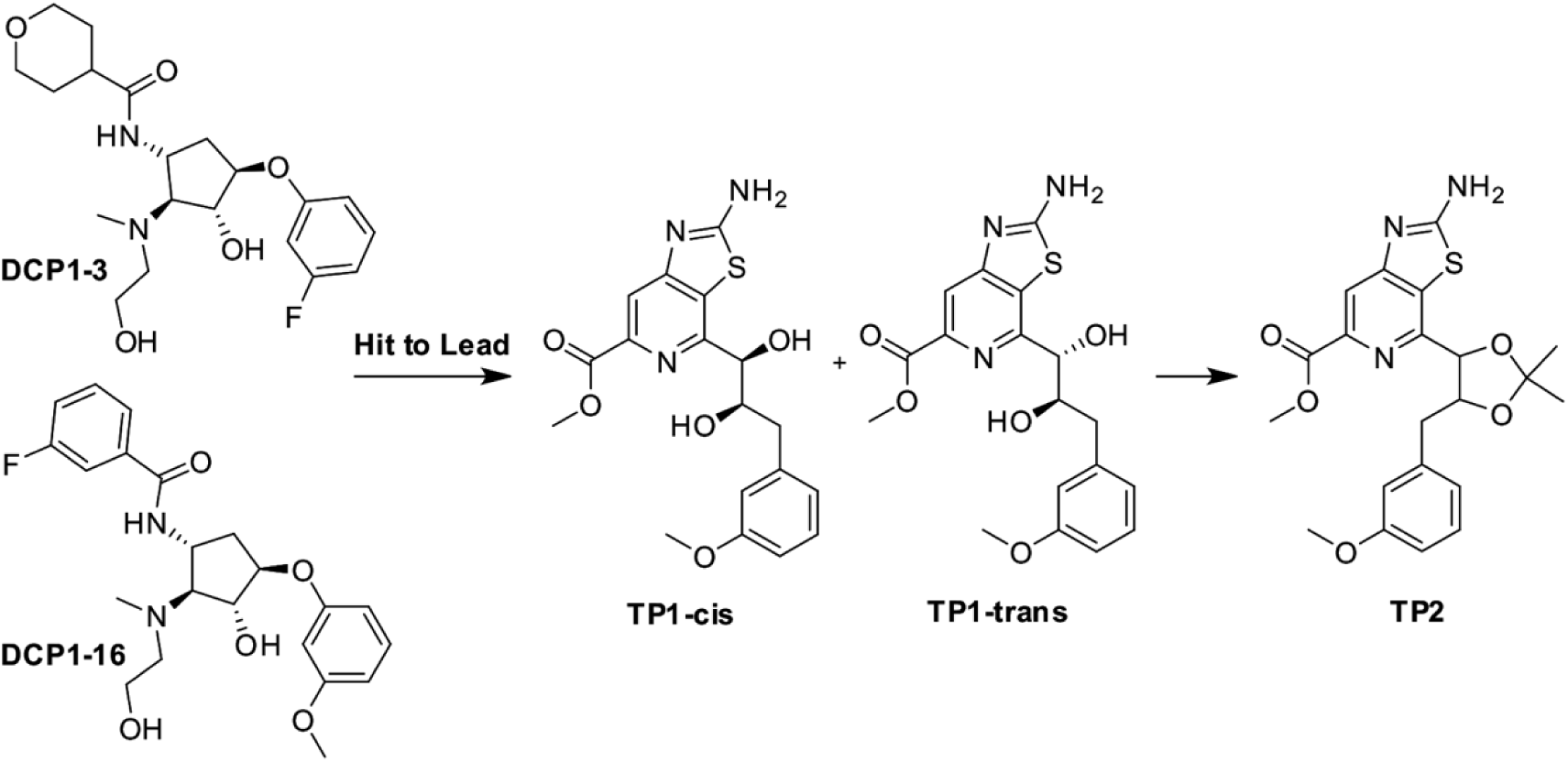
Hit-to-Lead Evolution of AT1R Allosteric Modulators. Structural refinement of initial non-drug-like hits DCP1-3 and DCP1-16 led to the design of the TP1 series, conserving the 3-methoxyphenyl aromatic vector and polar hydrogen-bonding network while replacing the cyclopentane core with a rigid thiazolo[4,5-c]pyridine scaffold. Synthesis yielded TP1-*cis* and TP1-*trans* diastereomers. Protection of the vicinal diol moiety yielded TP2, confirming the critical necessity of the free diol for biological efficacy (also see **Fig. S2 and Supplementary Methods**).

To progress from hit to lead, our goal was to develop ECL2 allosteric site targeted NAMs with <1 µM efficacy on signaling and the ability to lower blood pressure in AngII-infused mice. We pursued a structure-guided redesign rather than iterative substituent modification of the cyclopentane scaffold. Using molecular docking at the ECL2 allosteric site of the AT1R-AngII complex, we designed TP1, a compound built on a rigid thiazolo[4,5-c]pyridine core. TP1 conserves the two pharmacophoric elements essential for ECL2 allosteric site engagement, a polar hydrogen bonding hydroxyl motif and a hydrophobic methoxyphenyl aromatic vector, while eliminating the cyclopentane ring, the tetrahydropyranyl cap, the ethanolamine tail, and halogen substitutions present in DCP1-hits (**Fig. 1**). In place of the single hydroxyl found in DCP1-3 and DCP1-16, TP1 incorporates a vicinal diol linked to a meta-methoxybenzyl group, positioned to engage the same polar and hydrophobic sub-pockets occupied by the parent hits while introducing an additional hydrogen bond donor at this site.

To computationally test the functional importance of this diol prior to synthesis, we designed TP2 (**Fig. 1**), a masked analogue in which the vicinal diol is protected as a cyclic acetonide, eliminating its capacity to donate hydrogen bonds while preserving the remainder of the thiazolopyridine scaffold. Comparative Glide docking of the two designs predicted a substantially reduced binding score for TP2 relative to TP1 (-7.54 kcal/mol versus -9.51 kcal/mol), providing a basis for prioritizing TP1 as our lead candidate.

We examined the docked AT1R-TP1 complex to visualize the structural basis for activity. TP1 occupies a discrete pocket on the extracellular surface of AT1R, spatially distinct from the orthosteric AngII binding site (**Fig. 2A**). Within this pocket, the TP1 vicinal diol forms a direct hydrogen bonding network with residues H166, F170, H183, E185, N188 and T190 (**Fig. 2B and Figure S1**), consistent with the pharmacophore rationale underlying the design. In contrast, acetonide protection of the diol in TP2 redirects binding contacts away from this core network toward N168, H166, H183, and the peripheral, N-terminal residues K12 and Q15 (**Fig. 2C**). This shift in binding pose parallels the reduced Glide docking score of TP2 relative to TP1 (**Fig. S1**), suggesting reduced functional efficacy of TP2 in subsequent assays. Thus, our structure-guided hit-to-lead design and optimization exercise resulted in a novel chemotype, TP1, which preserves the key pharmacophoric elements required for receptor engagement while offering a synthetically feasible and computationally validated lead for further development.

**Figure 2.**
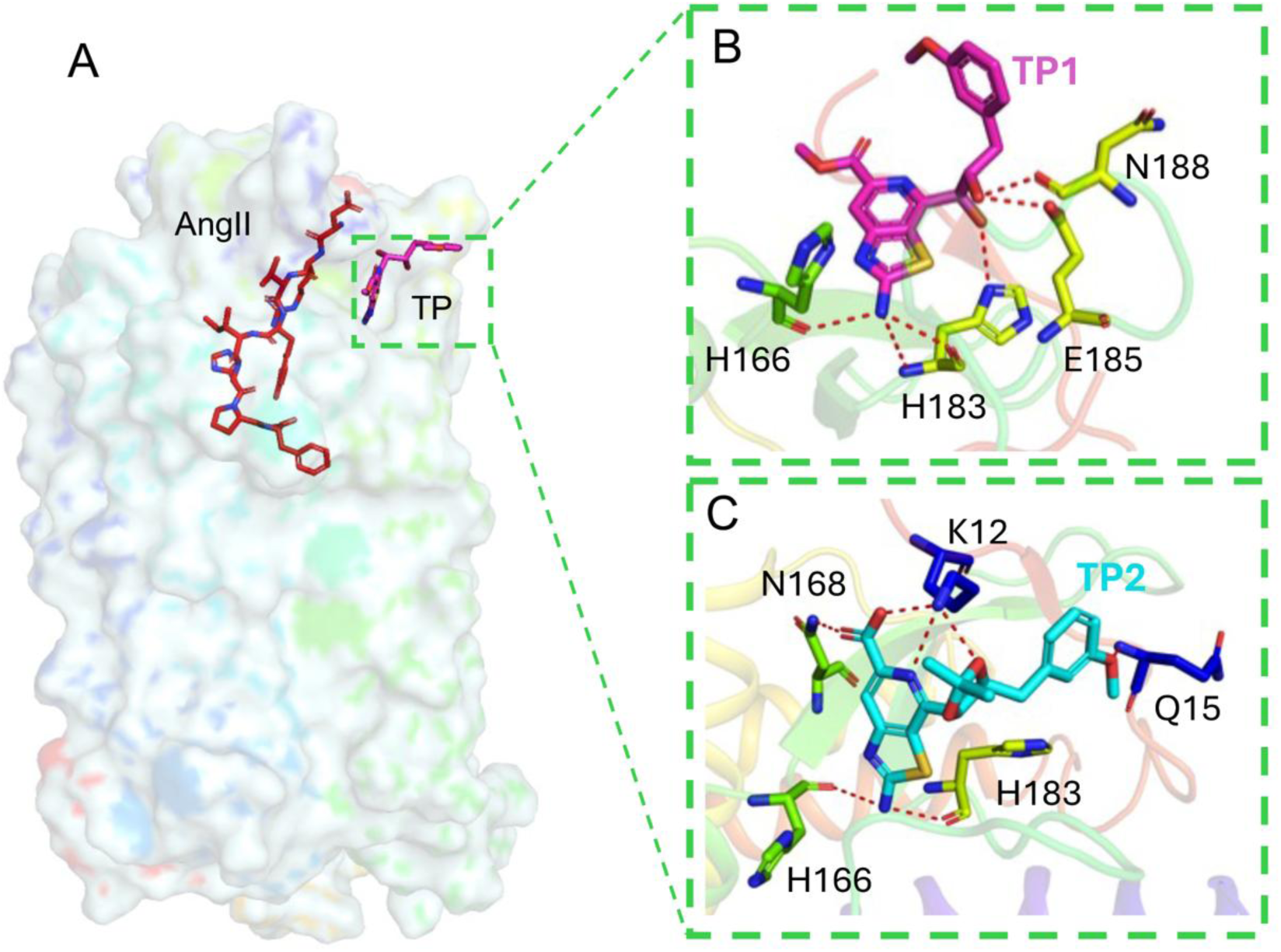
Molecular Docking Reveals Distinct Allosteric Engagement of TP1 and Disruption of the Binding Mode in TP2. (A) Electrostatic surface representation of AT1R showing the co-bound orthosteric agonist AngII (red sticks) and the allosteric ligand TP1 (magenta sticks, boxed region), illustrating the spatial separation between the orthosteric AngII-binding site and the distal allosteric pocket. (B) Zoomed view of the TP1 binding pose, showing key hydrogen bonding contacts (red dashed lines) between the TP1 vicinal diol and residues H166, H183, E185, and N188 within the allosteric pocket. (C) Zoomed view of the TP2 binding pose, showing that acetonide protection of the vicinal diol redirects binding contacts toward N168, H166, H183, and the N-terminal residues K12 and Q15, disrupting the hydrogen bonding network observed for TP1 (also see **Fig. S1**).

### Synthetic Creation of TP1 and TP2

Next, we synthesized both designed compounds. TP1 was obtained as a mixture of cis- and trans-diastereomers (TP1-cis and TP1-trans) at the diol stereocenter. TP2 was synthesized as the corresponding acetonide protected analogue. Compounds were characterized by HPLC, LC-MS 1H-NMR and 1H-NMR.D2O to confirm >97% purity and isomeric identity prior to functional evaluation. The synthetic route to TP1 and TP2, proceeding through a functionalized pyridine intermediate and culminating in the diol-bearing and acetonide-protected final compounds, is detailed in **Fig. S2 and Supplementary Methods**.

### Calcium Signaling Studies Reveal Negative Allosteric Modulation by TP1

Measuring AngII-mediated intracellular Ca²⁺ release is a functional readout for activation of AT1R signaling. To assess the impact of the TP compounds, HEK293T cells stably overexpressing AT1R (HEK-AT1R) were stimulated with doses of AngII in the absence or presence of TP1 or TP2, and the Ca²⁺ response was measured.

Treatment with TP compounds without AngII did not produce a Ca²⁺-response in cells (**Fig. S3)**, ruling out potential intrinsic agonistic activity of TP compounds. AngII-induced calcium mobilization was differentially attenuated in the presence of TP compounds consistent with a functional NAM. A pronounced rightward shift in the Ca²⁺ response curve was observed in the presence of TP1, which retains an unmodified diol moiety. The shift produced corresponds to ∼4.5-fold increase in the EC₅₀ (**Fig. 3A, 3C**). In contrast, TP2 with sterically blocked diol functionality elicited a modest ∼2-fold shift (**Fig. 3B, 3C**). The observed divergence in functional impact stresses the importance of the free diol group for productive receptor engagement, and allosteric activity in agreement with docking simulation data.

**Figure 3.**
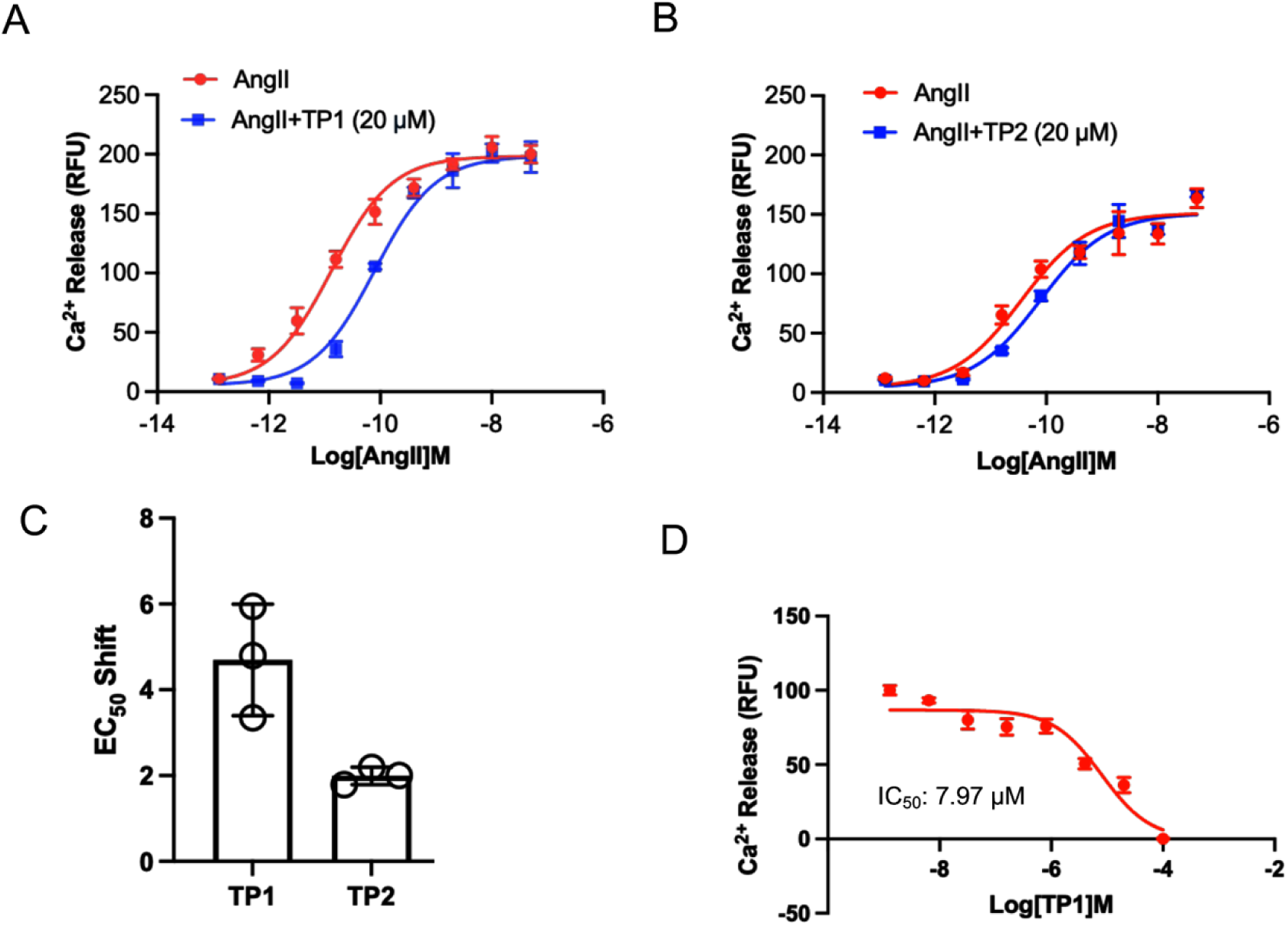
TP1, but not TP2, Shifts AngII-Induced Calcium Mobilization Rightward in HEK-AT1R Cells, Consistent with Negative Allosteric Modulation. **(A)** Concentration-response curves for AngII-induced intracellular Ca²⁺ release (measured as relative fluorescence units, RFU) in the absence or presence of TP1 (20 μM). Co-incubation with TP1 produced a clear rightward shift in the AngII concentration-response curve, indicative of negative allosteric modulation of AT1R. Data are mean ± SEM from n = 3 independent experiments. **(B)** Concentration-response curves for AngII-induced Ca²⁺ release in the absence or presence of TP2 (20 μM). In contrast to TP1, co-incubation with TP2 produced a comparatively modest shift in the AngII concentration-response curve. Data represents mean ± SEM from n = 3 independent experiments. **(C)** EC_50_ shift for TP1 and TP2, calculated as the foldchange in AngII EC_50_ in the presence of each compound relative to AngII alone, derived from the concentration-response curves in (A) and (B), quantitatively confirming a substantially greater rightward shift for TP1 than for TP2. **(D)** Concentration-response curve for TP1 inhibition of Ca²⁺ release in cells stimulated with AngII at its EC_20_ concentration, yielding an IC_50_ of 7.97 μM for TP1. TP2 produced no appreciable inhibition of AngII-induced Ca²⁺ release under these conditions and is therefore not shown. Data indicates mean ± SEM from n = 3 independent experiments (also see **Fig. S3**).

Effect of increasing doses of TP1 evaluated at submaximal EC_20_ concentration of AngII (0.025 nM) showed that NAM inhibitory potency yielding an IC₅₀ of approximately 7.97 μM (**Fig. 3D**). These findings demonstrate that TP1 is a NAM, selectively dampening AngII signaling, recapitulating the NAM property of precursor compounds, DCP1-3 and DCP1-16.

### TP1 Moderates AngII-Mediated Smooth Muscle Cell Migration and Wound Closure

We next assessed whether the NAM property of TP1 also regulates AngII-mediated vascular wall remodeling. AngII-induced scratch-wound closure in cultured mouse vascular aortic smooth muscle (MOVAS) cells are driven primarily by cell migration, and potentially by proliferation, making this assay useful for studying ligand effects on vascular wall repair and remodeling. We measured the effects of TP1 on MOVAS-AT1R cell migration using the scratch-wound assay, as described in our previous study^42^. AngII and Olmesartan were included as controls. Dynamic changes in scratch-gap width were measured at 8 hours, and total cell migration into the scratch area was measured at 24 hours of indicated treatments under serum-free conditions (**Fig. S4**).

The scratch-gap width decreased significantly following treatment with [Sar1] AngII (a stable analog of AngII) and [Sar1] AngII + TP1 compared with vehicle-treated cells and [Sar1] AngII + Olmesartan-treated cells. After 24 hours, cell migration into the scratch area was observed under both [Sar1] AngII + TP1 and [Sar1] AngII alone treatment but substantially increased compared with saline control. However, quantification showed that [Sar1] AngII + TP1 treatment allowed significant cell migration, although the magnitude was reduced compared with [Sar1] AngII alone. In contrast, cell migration following [Sar1] AngII + Olmesartan treatment was not significantly different from the saline control (**Fig. S4**). Together, these results suggest that TP1 moderates AngII-induced cell migration, and potentially proliferation, rather than completely blocking these effects. This partial regulation is consistent with an allosteric mechanism of action.

### Allosteric Modulation of Orthosteric Ligand Binding by TP1

To characterize the TP1-induced conformational effect on the orthosteric pocket, we performed ¹²⁵I-AngII saturation binding assays in the absence and presence of TP1. Co-incubation with TP1 produced a significant reduction in the B_max_ of ¹²⁵I-AngII binding to the orthosteric pocket (**Fig. 4A**). Nonlinear regression analysis of the binding isotherms revealed that decrease in B_max_ in the presence of TP1 is significant (p = 0.0129), whereas decrease in equilibrium dissociation constant (K_d_) was not significant (p = 0.2295) (**Fig. 4B-C**). This B_max_-selective effect, in the absence of a significant change in K_d_, indicates that TP1 likely reduces the number of AT1R binding sites competent to engage ¹²⁵I-AngII rather than altering the intrinsic affinity of the orthosteric pocket, which further supports negative allosteric modulation of AT1R by TP1.

**Figure 4.**
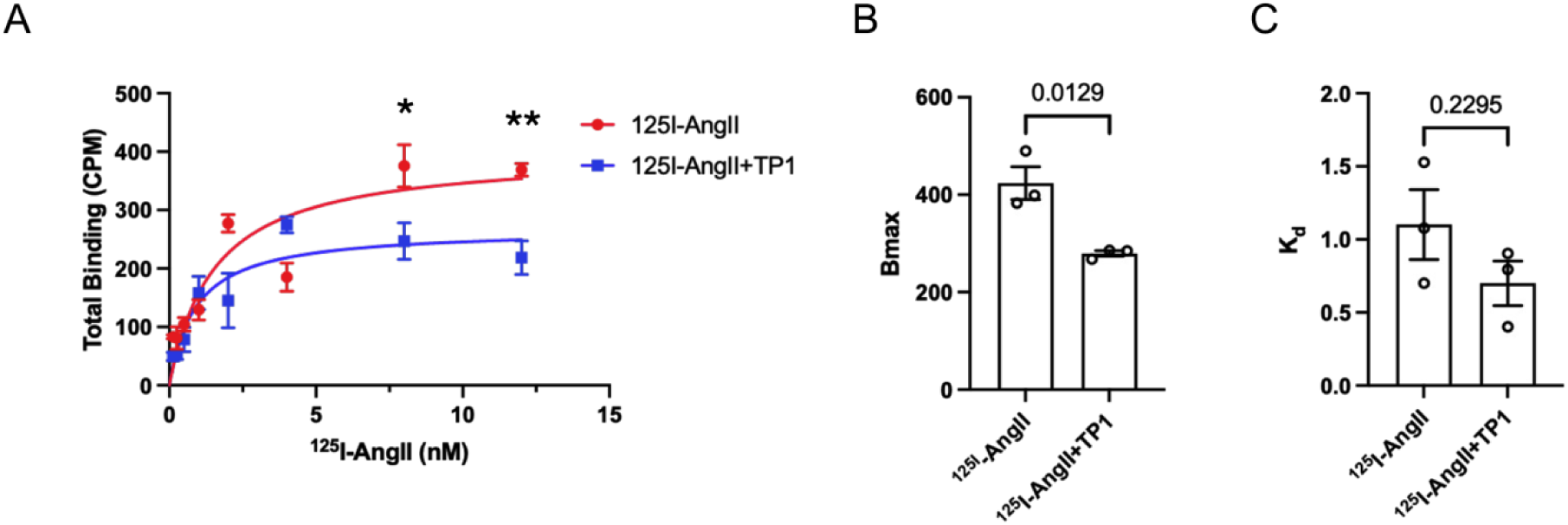
TP1 Reduces AT1R B_max_ Without Significantly Altering ¹²⁵I-AngII K_d_, Consistent with Negative Allosteric Modulation. **(A)** Saturation binding curves of ¹²⁵I-AngII in the absence or presence of TP1 (20 μM). Total binding was significantly reduced by TP1 co-incubation at higher ¹²⁵I-AngII concentrations (8 and 12 nM; *p* < 0.05 and *p* < 0.01, respectively). **(B)** B_max_ values derived from nonlinear regression of the saturation binding curves in (A), showing a significant reduction in the presence of TP1 (*p* = 0.0129).**(C)** K_d_ values derived from the same analysis, showing no significant difference in the presence of TP1 (*p* = 0.2295). Data are presented as mean ± SEM from n = 3 independent experiments.

### Mutagenesis Confirms TP1 Binding within the ECL2 Allosteric Pocket

To validate TP1 binding at the modeled ECL2-allosteric pocket, we performed computational alanine scanning on the docked AT1R-AngII-TP1 ternary complex (**Fig. 5A**). All residues lining the ECL2-allosteric pocket were targeted, instead of restricting to residues predicted to form direct hydrogen bonds in the docking model. Binding free energy change (Δaffinity = Affinity_mutant_ − ΔAffinity_wild-type_) was calculated for each substitution. Six Ala substitutions, H166A, F170A, N188A, H183A, E185A, and T190A, predicted the largest destabilization of TP1 binding (**Fig. 5B**). The docking model highlights H166, H183, E185, and T190 as residues directly engaged in hydrogen bonding with the TP1 (**Fig. 2B**), whereas F170 and Q187 contribute to the overall architecture of the binding pocket without making primary polar contacts in the docking pose (**Fig. S1**).

**Figure 5.**
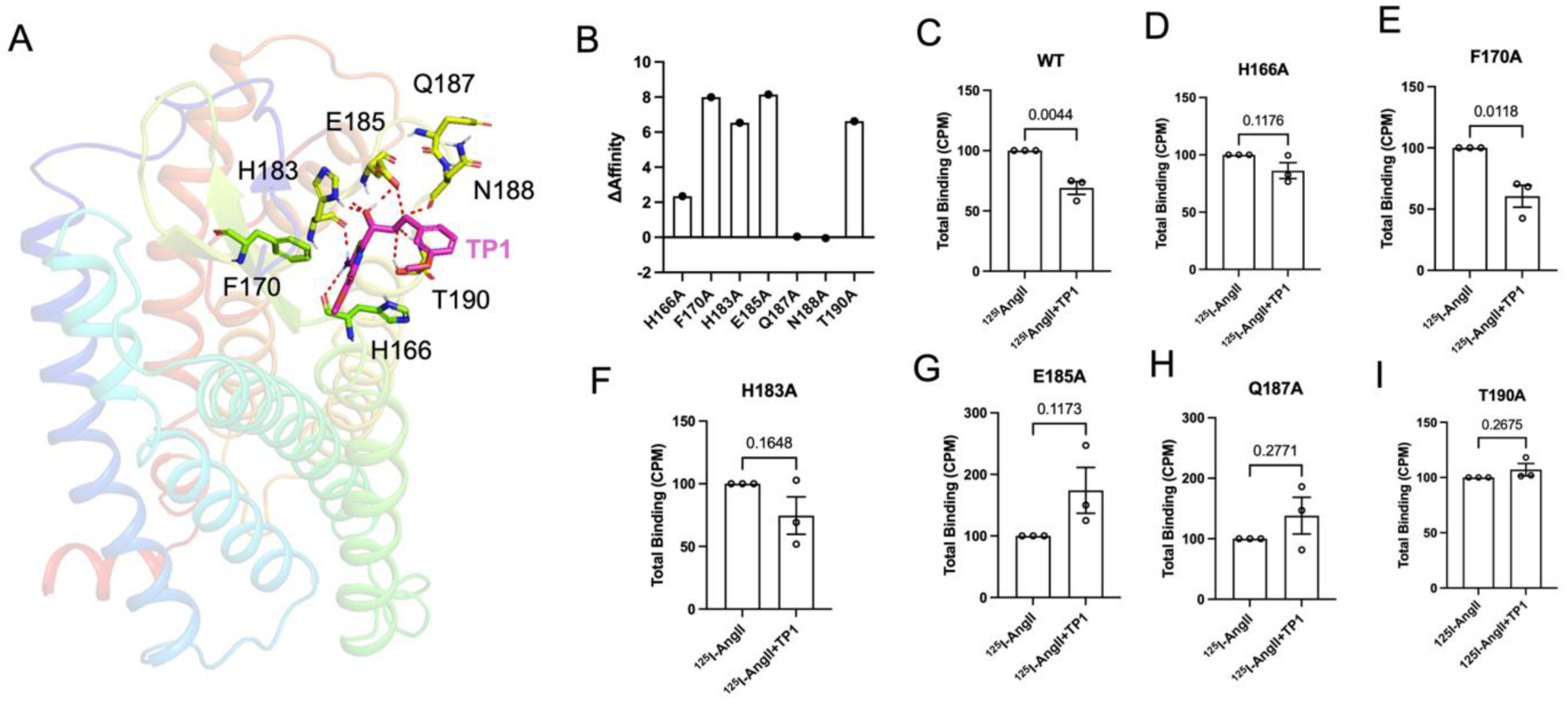
Computational and Experimental Alanine Scanning Confirms TP1 Binding Within the ECL2 Allosteric Pocket. **(A)** Docked structure of the AT1R-AngII-TP1 ternary complex showing residues lining the predicted TP1 allosteric binding pocket (H166, F170, H183, E185, Q187, T190). **(B)** Computational alanine scanning of the TP1 binding pocket. Bar graph shows the predicted change in binding free energy (ΔAffinity = ΔAffinity_mutant_ − ΔAffinity_wild-type_) upon in silico alanine substitution at each residue; higher ΔAffinity values indicate greater predicted destabilization of TP1 binding. **(C-I)** Experimental validation by ¹²⁵I-AngII saturation binding in WT and alanine mutant AT1R constructs (H166A, H183A, E185A, Q187A, T190A) transiently expressed in HEK293T cells, performed in the absence or presence of TP1. TP1 significantly reduced total ¹²⁵I-AngII binding in WT AT1R but not in any of the five alanine mutants, indicating loss of TP1-mediated negative allosteric activity upon mutation of predicted contact residues. Data shows mean ± SEM from n = 3 independent experiments; statistical comparisons by unpaired *t*-test.

To experimentally validate the computational alanine scanning findings, we generated six-point mutants of AT1R (H166A, F170, H183A, E185A, Q187A, and T190A) by site-directed mutagenesis and transiently expressed each mutant in HEK293T cells. An N188A mutant was also attempted but exhibited poor expression and was therefore excluded from subsequent binding analysis. Membrane preparations from WT and mutant transfected cells were subjected to single point binding analysis using 12 nM ¹²⁵I-AngII, the concentration at which the largest statistically significant difference in total binding was observed (**Fig. 4**), in the absence or presence of TP1 (20 μM). A significant reduction in total ¹²⁵I-AngII binding was observed in the WT AT1R in presence of TP1 as anticipated. In contrast, TP1-mediated inhibitory effects were lost across all six Ala substituted mutants, indicating that mutation of any single residue within the ECL2-allosteric pocket is sufficient to abrogate interaction with TP1 (**Fig. 5C-I**). Together, our computational and experimental mutagenesis data demonstrate that TP1 engages residues H166, F170, H183, E185, Q187, and T190 within ECL2 allosteric pocket to exert its NAM effect on AngII binding (**Fig. 2B; Fig. S1**).

### Vasomotor Function Analysis Confirms TP1 Activity in Artery Explants

To determine whether allosteric inhibition of AngII interaction with AT1R by TP1 translates into a functional physiological outcome at the tissue level, we assessed AngII-induced vasoconstriction using wire myography in isolated arterial segments. We selected the C57BL6 mice iliac artery for this analysis based on the expression level of AT1R and robust contractile responsiveness to AngII, as established in our prior studies^17^. Vessel rings mounted in the wire myography chamber maintained at resting tension were pre-incubated with either vehicle (physiological salt solution, PSS) or TP1 (20 μM) for 5 minutes prior to AngII (1 nM) stimulation and change in contractile responses were normalized to maximal contractile capacity produced by KPSS (high-potassium PSS) set as 100% to control for inter-vessel variability.

Treatment of PSS pre-treated iliac artery explants with TP1 alone did not produce any contractile response and/or affect vessel viability. Treatment with AngII elicited a robust vasoconstrictive response, whereas pre-treatment with TP1 significantly attenuated AngII-induced contraction (n = 8 vessels from N = 4 animals; p = 0.0362) (**Fig. 6**). This *ex vivo* functional inhibition of AngII-mediated vasoconstriction is consistent with our radioligand binding and calcium mobilization data and demonstrates that allosteric antagonism of TP1 at AT1R translates into functional suppression of AngII-driven vascular contraction.

**Figure 6.**
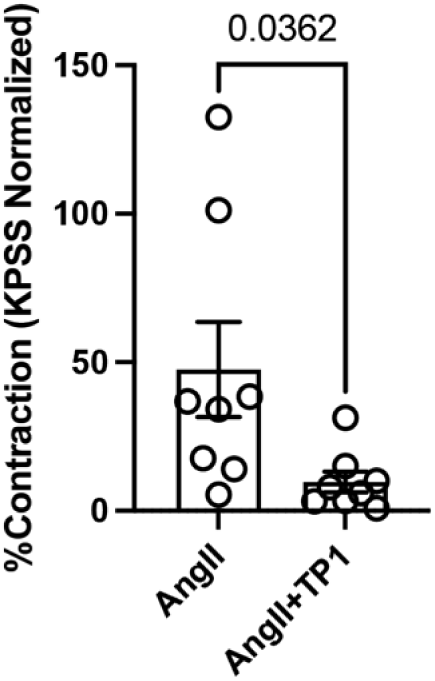
TP1 Attenuates AngII-Induced Vasoconstriction Ex Vivo. Wire myography analysis of AngII-induced contraction in isolated iliac artery from C57/BL6 mice segments pre-incubated with vehicle (PSS) or TP1 (20 μM) for 5 minutes prior to AngII (1 nM) challenge. Contractile responses are expressed as percent contraction normalized to KPSS-induced maximal contraction (%KPSS). TP1 pre-incubation significantly attenuated AngII-induced vasoconstriction relative to vehicle (*p* = 0.0362). Data are presented as mean ± SEM from n = 8 vessel segments from N = 4 animals per group.

### TP1 Significantly Attenuates AngII-Induced Hypertension *In Vivo*

To evaluate whether allosteric antagonism by TP1 translates into blood pressure (BP) lowering *in vivo*, we performed acute BP recordings using implantable radiotelemetry in conscious, freely moving C57BL6 mice. Baseline systolic and diastolic blood pressure (SBP and DBP) were recorded for 30 minutes prior to intraperitoneal (IP) administration of physiological salt solution (PSS, vehicle), AngII alone, AngII plus the AT1R-selective ARB Olmesartan, or AngII plus TP1, with all groups (n = 3 mice per group) receiving identical injection volumes to control for BP changes attributable to increased body fluid volume.

Peak elevation of SBP and DBP occurred at approximately 40 minutes, corresponding to 10 minutes post-injection. BP was recorded for 40 minutes following injection (**Fig. 7A, C**). No significant differences were observed upon administration of AngII alone, AngII+TP1 and AngII+Olmesartan during the rapid BP rise phase, but increases were significantly larger compared to a small change in BP observed in vehicle alone group. The AngII+Olmesartan group showed rapid decline in BP following the peak consistent with established blockade of AT1R by ARBs. The AngII+TP1 group in contrast showed a diminished decline over the same period, which suggests a moderate effect of TP1 on AngII induced BP increase.

**Figure 7.**
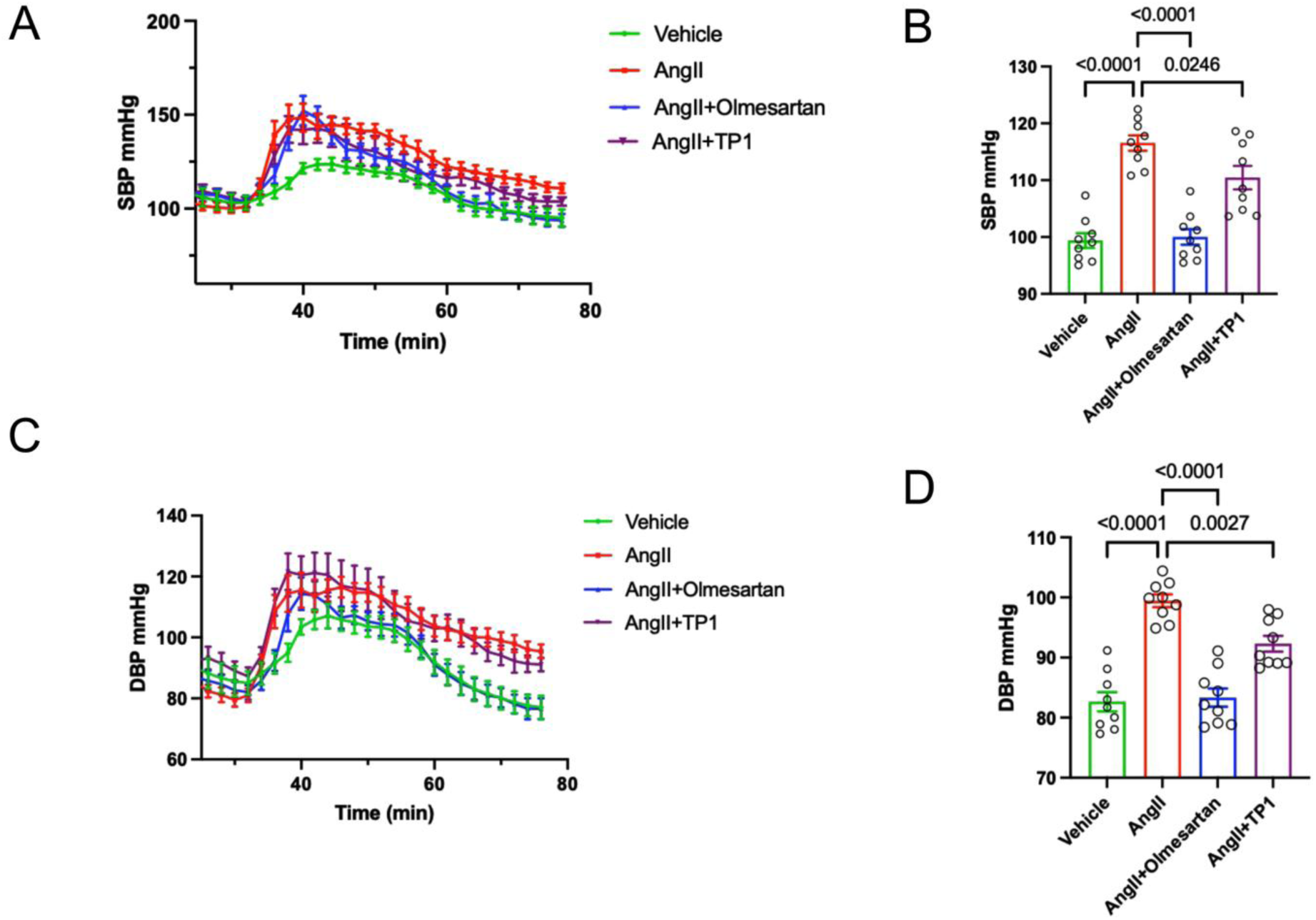
TP1 Attenuates AngII-Induced Hypertensive Responses In Vivo. Acute blood pressure responses were measured by radiotelemetry in conscious, freely moving mice following intraperitoneal infusion of vehicle, AngII alone, AngII plus Olmesartan, or AngII plus TP1, with all infusions initiated at the same time point. (A) Systolic blood pressure (SBP) time course. (B) Mean SBP averaged over the final 18 minutes of the recording period for each treatment group. (C) Diastolic blood pressure (DBP) time course for the same treatment groups. (D) Mean DBP averaged over the final 18 minutes of the recording period, calculated as in (B). All groups (n = 3 per group) received equal injection volumes on 3 days with 24 hrs. allowed for clearance between treatment. Data are presented as mean ± SEM. Statistical comparisons were performed by one-way ANOVA followed by Dunnett’s multiple comparisons test, with the AngII group set as the control column. Adjusted p values are shown above each comparison.

We quantified the change in BP from the peak response to the end of the recording period. Overall differences between groups were assessed using one-way ANOVA followed by Dunnett’s multiple-comparisons test, with AngII-treated mice used as the control group. During the decline phase, the reduction in SBP was significantly smaller in the AngII group than in both the AngII+Olmesartan group (mean difference = 16.52 mmHg; 95% CI: 11.11-21.93; p < 0.0001) and the AngII+TP1 group (mean difference = 6.09 mmHg; 95% CI: 0.67-11.50; p = 0.0246) (Fig. 7A, B). Comparing mean difference of SBP decline indicates that TP1 produced a smaller reduction in SBP than Olmesartan, highlighting the functional difference between inverse agonism and negative allosteric modulation (NAM). Decline of DBP showed a similar pattern. The change in DBP during the decline phase was significantly lower in the vehicle group (mean difference = 16.80 mmHg; 95% CI: 11.95-21.65; p < 0.0001) and the AngII+Olmesartan group (mean difference = 16.13 mmHg; 95% CI: 11.28-20.97; p < 0.0001) compared with the AngII group. The AngII+TP1 group showed a significantly lower change in DBP (mean difference = 7.15 mmHg; 95% CI: 2.30-11.99; p = 0.0027) (**Fig. 7C, D**). Overall, these results show that TP1, like Olmesartan, accelerates the resetting of AngII-induced BP elevation. However, this effect was stronger with Olmesartan than with TP1. Together, these findings demonstrate that TP1 significantly improves the resolution of AngII-induced hypertension in vivo. This provides functional evidence that TP1 acts as a NAM of AT1R and supports its potential as an antihypertensive agent in vivo.

### TP1 Blocks Binding of Pathogenic AT1R Autoantibodies from Patients with Preeclampsia and Dilated Cardiomyopathy

Pathogenic AT1R-AAbs have been implicated in the pathophysiology of both PE and DCM, where they engage ECL2 to drive sustained AT1R activation. To determine whether TP1 can interfere with this pathogenic mode of receptor engagement, we obtained serum samples from patients with DCM (n = 43) and PE (n = 84) and screened them by cell-based ELISA (cELISA) to identify samples with high AT1R-AAb titers. AT1R-AAb were present in >80% of patient sera in both DCM and PE cohorts (**Fig. S5**) From each cohort, we selected the three highest titer samples for further characterization.

We first confirmed the epitope specificity of these autoantibodies using a peptide competition binding assay. Serum samples from three high-titer DCM patients (DCM1168, DCM1177, DCM1197) and three high-titer PE patients (PE13, PE107, PE288) were incubated with HEK-AT1R cells in the absence or presence of a competing peptide corresponding to the ECL2 of AT1R. Pre-incubation with the ECL2 peptide significantly reduced antibody binding across all six samples tested, in both the DCM cohort (DCM1168, p = 0.005; DCM1177, p = 0.002; DCM1197, p = 0.03) and the PE cohort (PE13, p = 0.003; PE107, p = 0.02; PE288, p = 0.005). The presence of pathogenic autoantibody directed against the ECL2 epitope of AT1R was confirmed in both disease states (**Fig. S6**), which demonstrates a shared epitope specificity for AT1R-AAbs across two mechanistically distinct disease states.

We next tested whether TP1 could disrupt the interaction between AT1R and these antibodies. TP1 dose-dependently reduced binding of AT1R-AAb in serum samples from both DCM (IC_50_ = 0.85 µM) and PE (IC_50_ = 1.1 µM) patients (**Fig. 8A**). We further tested whether TP1 inhibitory activity extends beyond patient-derived polyclonal antibodies to a defined AT1R-ECL2 epitope directed monoclonal antibody previously reported by our group^17^. Dose-dependent inhibition of monoclonal antibody binding by TP1, with an IC_50_ of 0.1 μM was observed (**Fig. 8B**). Together, these findings establish that TP1 can inhibit not only patient-derived polyclonal AT1R-AAb but also a defined monoclonal antibody targeting the ECL2 epitope.

**Figure 8.**
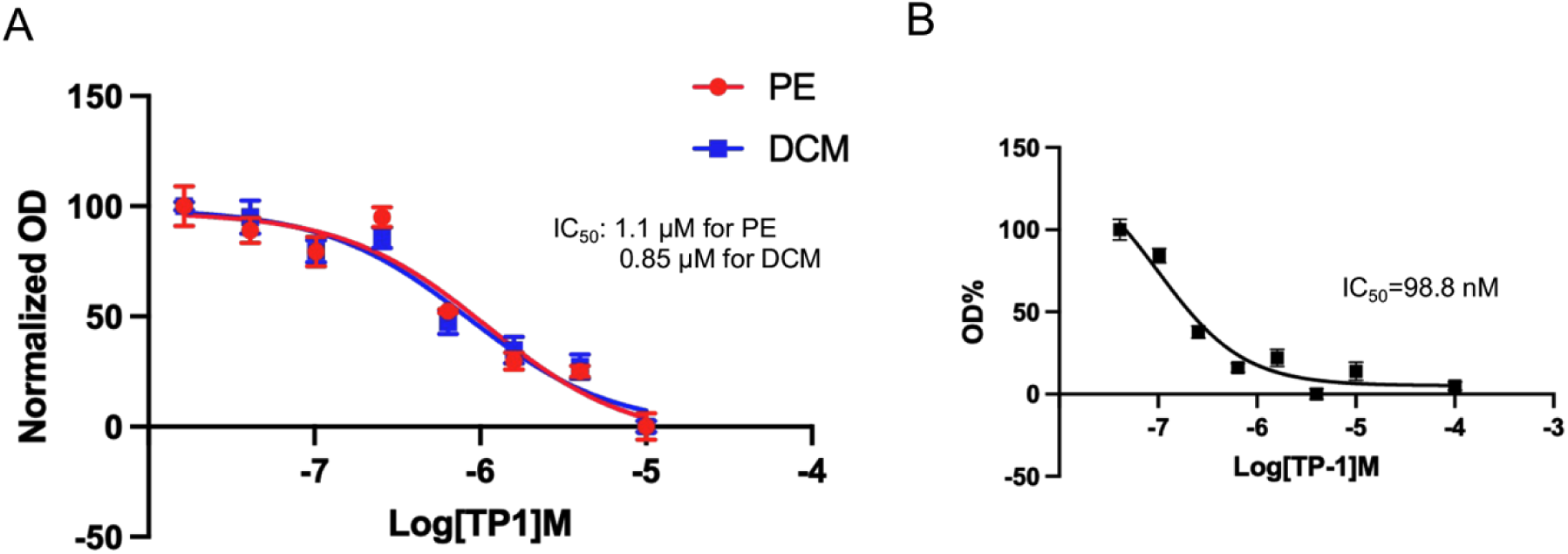
TP1 Dose-Dependently Inhibits Binding of Pathogenic AT1R Autoantibodies and a Monoclonal Anti-AT1R ECL2 Antibody. (A) Concentration-response curves showing TP1-mediated inhibition of AT1R autoantibody (AT1R-AA) binding, measured by cell-based ELISA, using serum samples from patients with preeclampsia (PE, red) and dilated cardiomyopathy (DCM, blue). TP1 produced comparable dose-dependent inhibition of autoantibody binding in both patient cohorts, with an IC_50_ of 1.1 µM for PE and 0.85 µM for DCM. (B) Concentration-response curve showing TP1-mediated inhibition of binding of a monoclonal antibody directed against the ECL2 of AT1R, previously reported by our group, to AT1R-expressing cells fixed with 4% paraformaldehyde, measured by cell-based ELISA. TP1 produced dose-dependent inhibition of monoclonal antibody binding, with an IC_50_ of 98.8 nM. (also see **Fig. S5 and S6**)

### TP1 acts as an AT1R-selective negative allosteric modulator with no significant effect on AT2R

Given that AT1R and AT2R share considerable structural homology despite modest sequence identity (∼30%), we next assessed whether allosteric action of TP1 is selective for AT1R or extends to AT2R, using a reciprocal ¹²⁵I-AngII binding analysis approach. Membranes prepared from HEK-AT1R, and HEK-AT2R cells were used. To exclude nonspecific reduction of bound ¹²⁵I-AngII, we included Olmesartan and PD123,319, well characterized selective antagonists of AT1R and AT2R respectively.

In the HEK-AT1R membranes, TP1 significantly reduced ¹²⁵I-AngII binding relative to the ¹²⁵I-AngII alone control (p < 0.0001). The AT1R specific Olmesartan (p < 0.0001) caused reduction of ¹²⁵I-AngII binding as anticipated. In contrast, PD123,319, the AT2R selective antagonist, had no significant effect on ¹²⁵I-AngII binding to AT1R (p = 0.2074) (**Fig. 9A**). In the reciprocal experiment, ¹²⁵I-AngII binding to HEK-AT2R membrane was not significantly reduced by TP1 (p = 0.0768) and Olmesartan (p = 0.2327) (**Fig. 9B**). The AT2R specific PD123,319 significantly reduced ¹²⁵I-AngII binding to AT2R (p < 0.0001), as expected for an AT2R selective antagonist. Unlabeled AngII displaced ¹²⁵I-AngII binding to both AT1R and AT2R through orthosteric competition (p < 0.0001) as anticipated. These reciprocal binding data demonstrate that TP1 selectively inhibits AngII binding at AT1R without appreciable cross reactivity at AT2R, supporting its pharmacological profile as a clean, subtype selective allosteric modulator suitable for further therapeutic development.

**Figure 9.**
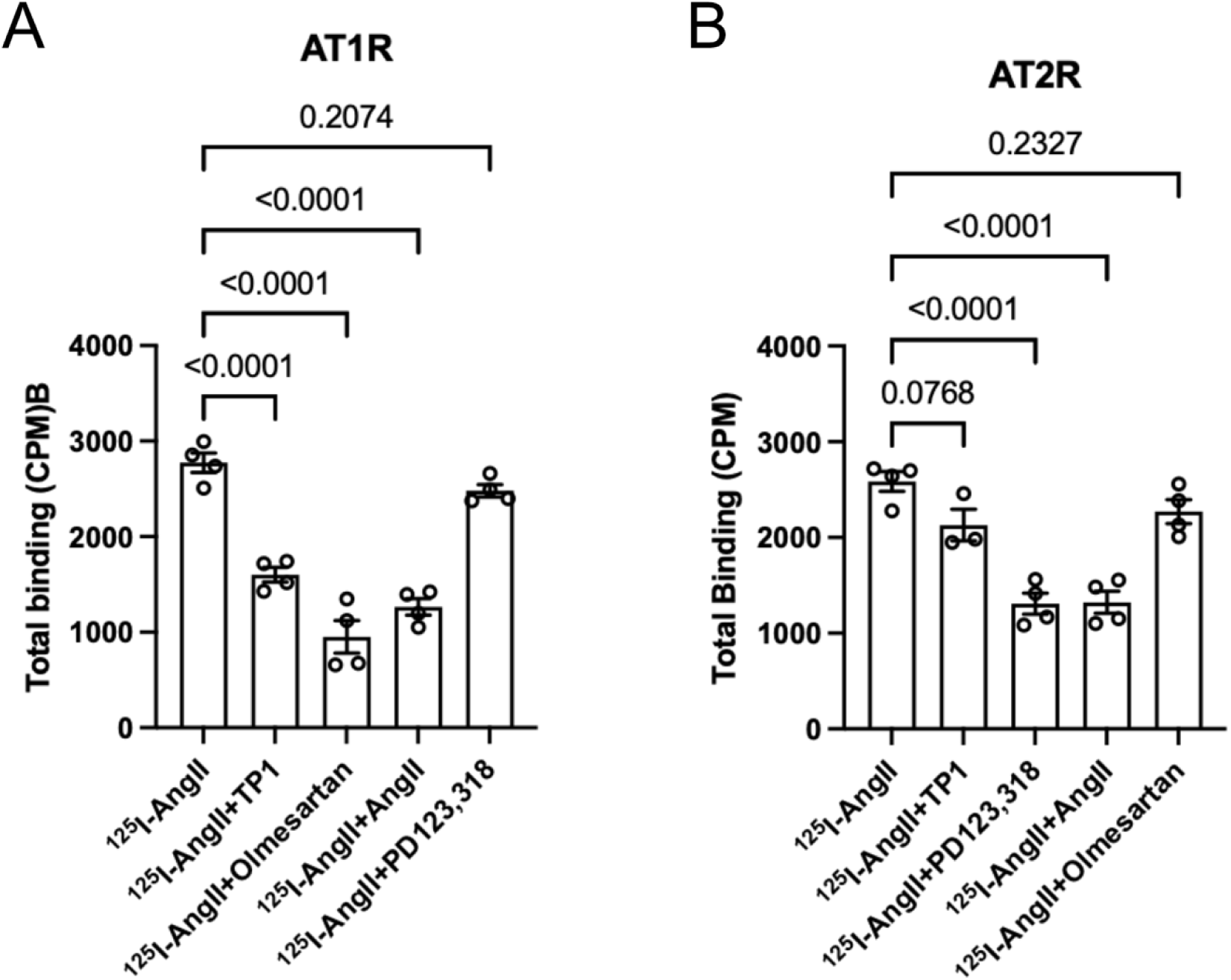
TP1 Selectively Inhibits AngII Binding to AT1R Over AT2R. Radioligand binding assays were performed using ¹²⁵I-AngII in membranes prepared from cells expressing AT1R (left) or AT2R (right). **(Left)** In AT1R expressing membranes, total ¹²⁵I-AngII binding was significantly reduced by TP1, the AT1R selective antagonist olmesartan, and unlabeled AngII, whereas the AT2R selective antagonist PD123,319 had no effect, confirming assay specificity for AT1R. **(Right)** In AT2R expressing membranes, total ¹²⁵I-AngII binding was significantly reduced by PD123,319 and unlabeled AngII, whereas olmesartan had no effect. TP1 similarly failed to significantly reduce ¹²⁵I-AngII binding at AT2R, paralleling the lack of effect observed with the AT1R selective control olmesartan. Data shown are from a representative experiment (n = 4 technical replicates per condition), independently repeated three times with consistent results. Statistical comparisons were performed by one-way ANOVA with post hoc multiple comparisons testing; adjusted p values are shown above each comparison.

### TP1 Does Not Exhibit Cytotoxicity in Cultured Cells

To evaluate whether TP1 exerts any nonspecific cytotoxic liability, we assessed viability and toxicity in HEK-AT1R cells using the CellTiter-Blue Cell Viability Assay and the CellTox Green Cytotoxicity Assay. Cells were treated with vehicle, AngII (20 μM), olmesartan (20 μM), or TP1 at concentrations 0.2, 2.0 and 20 μM. At 72 hours, cell viability was measured as resorufin fluorescence generated by metabolically active cells was comparable across all treatment groups. Cytotoxicity was measured as fluorescence from the CellTox Green dye that selectively labels DNA in cells with compromised membrane integrity. (**Fig. S7**).

Viability in the TP1 treated cells was comparable to all treatment groups, including vehicle, AngII, Olmesartan, with no statistically significant differences observed by one-way ANOVA with Dunnett’s multiple comparisons test (p > 0.05 for all comparisons) (**Fig. S7A**). Cytotoxicity was low at background levels across vehicle, AngII, olmesartan, and all three TP1 concentrations. As expected, the lysis buffer positive control produced a robust and significant increase in cytotoxicity signal relative to vehicle (p < 0.0001), confirming that the assay was appropriately sensitive to detect genuine loss of membrane integrity when present (**Fig. S7B**). Together, these results demonstrate that TP1 does not impair cell viability or induce cell death at concentration substantially exceeding its pharmacologically active concentration at AT1R, indicating that inhibition of AngII effects by TP1 are not confounded by general cytotoxicity.

### Integrated Pharmacokinetic Characterization Supports the Acute Dosing Paradigm Used for In Vivo Efficacy Studies

To contextualize TP1’s in vivo efficacy and guide its future optimization, we characterized its pharmacokinetic (PK) and physicochemical properties using an integrated in vitro-in vivo pharmacology framework, encompassing kinetic aqueous solubility, Caco-2 cell permeability, liver microsomal metabolic stability, plasma protein binding, and single-dose PK in rats (**Table 1**).

**Table 1.** Integrated pharmacokinetic and physicochemical profile of TP1. Full parameter sets, reference compound data, and individual replicate values are provided in SI Appendix, **Tables S1-S5**.

| Property | Assay | TP1 Value | Interpretation |
| --- | --- | --- | --- |
| Kinetic solubility (PBS, pH 7.4) | Shake-flask, LC-MS/MS | 82.8 ± 18.7 µM (of 100 µM nominal) | Good |
| Caco-2 permeability (Papp A→B) | Bidirectional transport | 1.43 × 10<sup>-6</sup> cm/s | Low |
| Caco-2 efflux ratio | Bidirectional transport | 12.9 | Potential P-gp substrate |
| Microsomal T<sub>1/2</sub> (rat / human) | Liver microsomes + NADPH | 15.0 / 25.9 min | Moderate-high clearance |
| Hepatic extraction ratio, Eh (rat / human) | Well-stirred liver model | 66.1% / 74.9% | Moderate-high clearance |
| Plasma protein binding (human) | Equilibrium dialysis | 88.6% bound (Fu = 11.4%) | Moderate |
| IV PK (2 mg/kg, rat) | Noncompartmental analysis | T<sub>1/2</sub> = 0.47 h; Cl = 460.2 mL/h/kg | High clearance |
| Oral bioavailability (10 mg/kg, rat) | Noncompartmental analysis | F = 15.3% | Low |

TP1 showed good aqueous solubility in PBS at physiological pH (82.8 ± 18.7 μM at a 100 μM nominal concentration), indicating that solubility is unlikely to limit its in vitro potency or in vivo formulation. In Caco-2 cell monolayers, however, TP1 displayed low apical-to-basolateral passive permeability (Papp = 1.43 × 10⁻⁶ cm/s) together with a high efflux ratio (12.9), identifying TP1 as a potential P-glycoprotein (Pgp) substrate and suggesting that both limited passive diffusion and active efflux may constrain its intestinal absorption. Consistent with this observation, TP1 underwent moderate-to-high metabolic turnover in liver microsomes, with an in vitro half-life of 15.0 min in rat and 25.9 min in human microsomes and predicted hepatic extraction ratios of 66.1% and 74.9%, respectively, indicating substantial first-pass metabolism in addition to limited absorption. TP1 was moderately bound to human plasma protein (88.6% bound; Fu = 11.4%), supporting adequate free-drug availability at pharmacologically active concentrations.

The in vitro liabilities converged with TP1’s in vivo PK profile in rats. Following IV administration (2 mg/kg), TP1 was cleared rapidly (T_1/2_ = 0.47 ± 0.27 h; Cl = 460.2 ± 183.6 mL/h/kg) with a large volume of distribution (Vss = 6671 ± 3498 mL/kg), while oral administration (10 mg/kg) yielded low absolute bioavailability (F = 15.3 ± 1.9%) (**Fig. 10, Fig S8, Table 1**). Together, this integrated PK/PD profile, low oral absorption compounded by high systemic clearance, provided the pharmacological rationale for the acute, IV-dosing paradigm used in our in vivo efficacy studies (radiotelemetry blood pressure monitoring), rather than chronic oral dosing, and defines the principal pharmacokinetic liabilities to be addressed in future lead optimization of the TP1 series. Full assay-specific methods, individual compound-level data, and reference compound validation are provided in **Tables S1-S5, Fig. S8 and Supplementary Results.**

**Figure 10.**
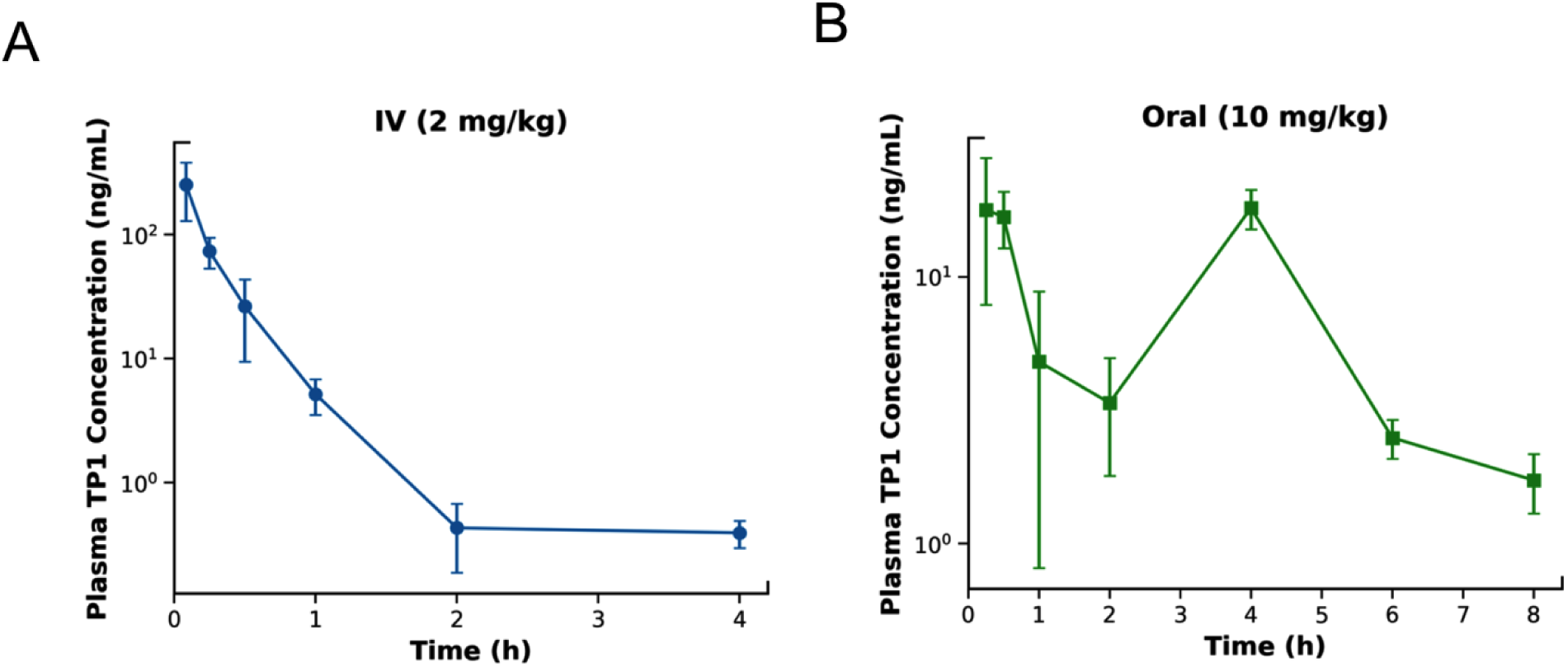
Pharmacokinetic Profile and Systemic Clearance of TP1 Following Intravenous and Oral Administration: TP1 shows rapid systemic clearance after IV dosing and low oral bioavailability in rats. Mean plasma concentration-time profiles of TP1 (log scale) following a single intravenous dose (2 mg/kg, left) or oral dose (10 mg/kg, right) in male Sprague-Dawley rats. Data are presented as mean ± SD (n = 3 animals per group). (also see **Fig. S8 and Supplementary Results**)

## Discussion

This study reports the hit-to-lead advancement of two previously identified compounds, DCP1-3 and DCP1-16, which we reported as potential NAMs of AT1R^17^. Using structure-guided virtual design and optimization, the novel lead compound TP1 was developed to target the cryptic allosteric site of AT1R, which also encompasses the epitope recognized by AT1R-AAbs. Synthesized TP1 was experimentally tested for its efficacy in cell signaling, vessel contraction, live-animal blood pressure and patient serum AT1R-AAb binding. TP1 provides four important features distinct from any AT1R ligands. First, it is a novel chemical and has no structural similarity to known AT1R ligands. Second, TP1 binds to an allosteric site on the extracellular region of AT1R not directly competing for the membrane embedded orthosteric site. Third, TP1 allosterically alters the accessibility of orthosteric pocket to the natural agonist AngII. Fourth, when TP1 binds to AT1R, it masks/disrupts the common epitope recognized by AT1R-AAbs found in autoimmune diseases. As a proof-of-concept, TP1 thus expands the pharmacological toolkit for selectively inhibiting AT1R signaling through its allosteric site. Our work also provides a foundation for developing a new generation of AT1R-targeted therapies that may offer advantages over currently available ARBs.

Functional characterization of TP1 directly confirmed key computational predictions. The hydrogen-bonding vicinal diol motif, which connects the hydrophobic methoxyphenyl group to the rigid thiazolo[4,5-c]pyridine core, is essential for engaging the ECL2 allosteric site, which comprises of hydrophilic residues His, Thr, Gln, and Glu. The comparable allosteric activity of TP1-cis and TP1-trans demonstrated that the stereochemical configuration of the vicinal diol does not determine activity, allowing both diastereomers to be carried forward. Furthermore, the computationally predicted loss of binding affinity for TP2 was consistent with its experimentally observed loss of functional efficacy. This agreement validates the docking model as a reliable predictor of structure–activity relationships within this series and identifies the vicinal diol as the central pharmacophore responsible for TP1 activity.

Our medicinal chemistry strategy developing TP1 overcame several practical limitations of the initial hits, DCP1-3 and DCP1-16, including their limited availability from a commercial screening library, which could not be readily replenished, and their multiple stereocenters, which complicated the attempts for independent synthesis and optimization.

Potency of TP1 improved compared with the original hits in the Ca²⁺ signaling assay. DCP1-3 and DCP1-16 showed no detectable NAM activity against the physiologically agonist, AngII and its more potent, stable analog [Sar1] AngII. They were therefore characterized using AngIV, which is a partial agonist of AT1R with approximately 100-fold lower affinity. Because allosteric effects are probe-dependent, a direct fold-potency comparison between TP1 and the original hits has not been tested using the same orthosteric probe. Nevertheless, the Ca²⁺ signaling and cell migration data demonstrate that TP1 is active against AngII and [Sar1] AngII, while the original hits were not, supporting a meaningful improvement in potency. The functional potency of TP1 against AngII-induced Ca²⁺ signaling (IC₅₀ ≈ 7.97 μM) is consistent with that reported for first-generation NAMs developed through structure-based drug discovery campaigns targeting other GPCRs. For example, mGlu5 NAMs have been reported with affinities ranging from 0.43 to 8.6 μM^43^. Thus, the potency of TP1 is within the range observed for early-stage GPCR NAMs and provides a reasonable starting point for future medicinal chemistry optimization to further improve potency.

As a NAM, TP1 alters AT1R conformation, reduces rather than completely suppress the binding of ¹²⁵I-AngII, Ca²⁺ signaling, cell migration (proliferation) and wound-healing, which may be therapeutically desirable. Chronic AT1R signaling is a well-established driver of NF-κB activation, cytokine production, and immune cell activation, independent of blood pressure lowering^44^. Thus, selectively attenuating pathological, inflammation-driving AT1R activity by TP1 while preserving sufficient receptor signaling for normal physiological function may be particularly relevant for chronic, low-grade inflammatory conditions. Direct comparison of negative allosteric modulation by TP1 with complete AT1R blockade by ARBs in relevant disease models will be needed to further verify this hypothesis.

Mechanistic insight for how TP1 may cause partial modulation is provided by the ¹²⁵I-AngII binding data. Classical negative allosteric cooperativity is expected to manifest as an increase in the K_d_ of the orthosteric radioligand, with B_max_ unchanged. Precedent for a B_max_-selective rather than K_d_-selective effect exists at other class A GPCRs; for instance, G protein coupling at the β2-adrenergic receptor alters the apparent B_max_ of orthosteric ligand binding by stabilizing a distinct receptor conformation that restricts access to and egress from the orthosteric pocket, independent of a classical affinity-based mechanism^45^. Thus, allosteric modulators are not obligated to follow a classical pattern and may or may not affect the K_d_ of the orthosteric radioligand^46^. Our data demonstrating the B_max_-selective effect for TP1 is consistent with stabilization of a subpopulation of AT1R in a conformation refractory to productive AngII engagement. Future biophysical studies are needed to confirm this hypothesis.

Reciprocal ¹²⁵I-AngII binding studies demonstrated that TP1 is selective for AT1R, with no appreciable cross-reactivity with AT2R. This selectivity is pharmacologically important because AT2R generally competes against AT1R signaling and promotes vasodilation, natriuresis, and anti-proliferative and anti-inflammatory effects, which are considered tissue and organ protective^47–49^. Blocking both AT2R and AT1R could interfere with these beneficial pathways. Since TP1 does not affect AT2R signaling, its potential as a selective NAM for further development is very promising.

The NAM activity of TP1 translated into a live-animal physiological effect. Our blood pressure measurement by radiotelemetry for assessing systolic and diastolic blood pressure in conscious rodents is a gold-standard method^50^. TP1 attenuated AngII-induced blood pressure elevation, with a time course like that of Olmesartan, a clinically approved ARB used as a control in the same experiment. This head-to-head comparison demonstrates that a physiologically meaningful antihypertensive effect can be produced from negative allosteric modulation of AT1R. Even though TP1 currently has unoptimized pharmacokinetic profile with a short half-life of less than one hour, it still produced a clear and statistically significant acute response. This suggests that TP1 was able to reach and affect its target, AT1R, enough to produce an effect on the whole animal. This provides an important insight on behalf of TP1 as an early-stage lead candidate suitable for further optimization of its pharmacokinetic properties.

Computational modeling and experimental mutagenesis demonstrated that TP1 directly binds to the same epitope on AT1R recognized by AT1R-AAbs, which was the main goal of our lead design campaign. cELISA results further confirmed that TP1 binds at the receptor–autoantibody interface by showing that it blocked AT1R-AAb binding from patients with PE and DCM, as well as binding of the surrogate monoclonal antibody. Therefore, TP1 may have the potential to block the interaction between AT1R and AT1R-AAbs across a broad range of diseases^21,51^. In pregnant women with PE, antihypertensive treatment with ARBs is contraindicated because of the risk of fetotoxicity^11,12^. PE is a well-established condition in which AT1R-AAbs are present in approximately 58%–90% of patients, depending on the study population^20,52^ but current treatment strategies do not directly target antibody-driven activation of AT1R. Instead, drug development efforts have focused on complex downstream effects of AT1R, including increased circulating soluble fms-like tyrosine kinase-1 (sFlt-1), impaired trophoblast invasion and spiral artery remodeling, and reduced placental perfusion and fetal growth^40,52^. The complex pathogenesis of PE can directly affect fetal development and may have further limited efforts to develop AT1R-targeted therapies for this condition. In transplant rejection and malignant hypertension, ARBs have been shown to partially reduce the harmful effects driven by AT1R-AAbs^27,53–55^. Although this partial effect may reduce the urgency to develop drugs that directly target the antibody-receptor interface, such as TP1, it suggests that our current understanding of the pathogenic mechanisms of AT1R-AAbs may be incomplete.

Our strategy to block AAb binding to AT1R could, in principle, prevents both the established agonistic AAb signaling and the potential effects of the antigen-AAb complex through the complement (C1q) and Fcγ receptor effector pathways in AT1R-AAb-mediated diseases. AT1R-AAbs identified in these diseases are predominantly IgG1 and IgG3 subclasses, which are particularly capable of activating the complement system and engaging Fcγ receptors on effector cells^20,51^. Whether this Fc-mediated effector component contributes to additional pathogenic signaling pathways has not been directly investigated for AT1R-AAbs. Importantly, ARBs block only the orthosteric AngII binding pocket and cannot prevent antibody binding to ECL2. Therefore, any residual Fc-mediated effects, if present, would remain unaddressed. TP1 is designed to address this therapeutic gap. To our knowledge, no previous small molecule exists that directly blocks AT1R-AAb binding while preserving the normal AngII-AT1R signaling pathway.

The preliminary pharmacokinetic (PK) profile of TP1 compares favorably with peptide-based strategies for blocking AT1R-AAbs. Unmodified linear peptides used in previous AT1R-AAb blocking studies are rapidly degraded in plasma, with approximately 80% degraded within 30 minutes^40,56^. A retro-inverso-stabilized peptide showed improved stability, with approximately 70% remaining intact after 6 hours^39,56,57^. In contrast, TP1 has a measurable plasma half-life (T_½_ = 0.47 h, IV) as a small molecule emerging directly from an initial hit-to-lead campaign, without any dedicated metabolic stabilization efforts. Although this half-life is not yet optimized, it provides a useful starting point for further development. Small-molecule scaffolds can be further optimized through conventional medicinal chemistry approaches, making TP1’s current PK profile better viewed as an encouraging early-stage result rather than a compound-specific limitation.

Creation of TP1 raises an important question; should it be viewed as an AT1R antagonist like traditional ARBs, or as a dedicated AT1R-AAb blocker? Both roles are possible. As an AT1R-AAb blocker, TP1 targets a mechanism that current therapies do not directly address, with relevance to preeclampsia, where ARBs cannot be used. As a negative allosteric modulator, TP1 only partially suppresses AngII signaling, potentially allowing normal physiological AT1R activity while reducing excessive signaling. This may be useful in conditions such as mild hypertension or chronic inflammation. Overall, TP1 provides a promising prototype for targeting autoantibody-driven GPCR activation, a mechanism that may extend beyond AT1R.

Our future studies will address several limitations which were beyond the scope of the present study. In vivo chronic dosing studies, ideally using an analogue of TP1 with improved-PK administered orally, will be required to establish durable antihypertensive efficacy and to evaluate whether sustained AT1R allosteric modulation produces adaptive changes in receptor expression or signaling not captured in an acute paradigm. Our PK and plasma protein binding characterization relied on modest sample sizes (n = 2-3 per condition) typical of early-stage DMPK screening, and larger confirmatory cohorts would strengthen confidence in these estimates. The mechanistic basis for TP1’s disruption of AT1R-AA engagement remains correlative rather than structurally confirmed. While our alanine scanning approach validated the residues lining the TP1 binding pocket, it does not establish the precise conformational states involved in allosteric inhibition of the receptor, a question that would benefit from complementary biophysical approaches in future work.

## Conclusion

This study establishes TP1 as a structure-guided lead compound that resolves the low potency and poor drug-like properties of the DCP1 hit series. Crucially, TP1 retains the defining feature of these earlier compounds: allosteric engagement of the ECL2 pocket of AT1R, which is spatially and functionally distinct from the orthosteric site targeted by conventional ARBs. We validated TP1’s efficacy as a negative allosteric modulator across *in vitro, ex vivo*, and *in vivo* models, demonstrating a measurable reduction in blood pressure elevation despite an unoptimized pharmacokinetic profile. Most notably, TP1 selectively blocks pathogenic AT1R-AAbs isolated from preeclampsia and dilated cardiomyopathy patients, a mechanism entirely inaccessible to orthosteric ARBs and for which no small-molecule inhibitor has previously been reported. Together, our findings provide proof of concept that allosteric targeting of the AT1R can simultaneously modulate physiological AngII signaling and neutralize pathogenic autoantibody activation. This positions TP1 as an important first step toward therapeutics for autoantibody-driven conditions such as pre-eclampsia, where conventional ARBs are contraindicated, and offers a broader framework for targeting autoantibody-mediated GPCR pathologies.

## Methods

### Molecular Docking

The starting coordinates of AT1R were retrieved from the Protein Data Bank (PDB ID: 6OS0; https://www.rcsb.org). Prior to docking, the crystallographic construct was modified to reflect the native receptor: the stabilizing nanobody Nb.AT110i1, used in the original structure to trap the active conformation via engagement of the intracellular transducer pocket, and the thermostabilized apocytochrome b562RIL (BRIL) fused into intracellular loop 3 (ICL3) to facilitate crystallization, were both removed, and the native AT1R ICL3 sequence was restored. The structure was then prepared for Glide docking calculations (Glide, Schrödinger, LLC)^58^ using the Protein Preparation Wizard, applying the default OPLS3 (Optimized Potentials for Liquid Simulations, Schrödinger, Inc.) force field energy minimization protocol^59^. Missing loop segments were modeled using Prime (Schrödinger, Inc.) during protein preparation^60^. Ligands were prepared using LigPrep (Schrödinger, LLC).

Ligand docking to the proposed allosteric site was performed using Schrödinger’s Induced Fit Docking (IFD) module. In the initial stage, ligands were docked into a rigid receptor model using Glide XP mode, with van der Waals (vdW) radii scaled to 0.5 for nonpolar atoms of both protein and ligand to accommodate minor steric clashes; the resulting ligand poses were retained for subsequent protein structural refinement. Constrained energy minimization was then performed using the OPLS3 force field with an implicit solvation model, relieving residual steric strain while maintaining overall protein conformation close to the original crystal structure. Prime (Schrödinger, LLC) was used to generate the corresponding induced-fit protein-ligand complexes. In the final stage, each ligand was redocked into the top 20 refined receptor structures using Glide XP and binding free energies for all resulting protein-ligand complexes were estimated using Prime/MM-GBSA, as described previously. Full details of the docking protocol have been reported in our previous work^8,17,61^.

### Calcium Assay

AngII-induced calcium mobilization was measured in HEK293T-AT1R stable cell lines generated by transfecting AT1R-negative HEK-293T cells, to model receptor-specific signaling in a widely used immortalized human cell model. The assay protocol was modified from previously published methods^17,42,62^ and utilized the Fluorescent Imaging Plate Reader (FLIPR) Calcium 5 Assay Kit (Molecular Devices). Cells (50,000 per well) were seeded in poly-D-lysine-coated, clear-bottom, black 96-well plates using a Cellometer T4 cell counter (Nexelcom Biosciences, Revvity, Chicago, IL, USA) and incubated for 24 h before serum starvation for 2 h to synchronize basal signaling. Cells were then loaded with Calcium 5 dye and kept for 0.5 h at 37°C in 5% CO₂ and 0.5 h at room temperature. Drug plates were separately prepared in a 96 well clear bottom with U-shape. Prepared in triplicate with dose-response of AngII and AngII plus 20 uM of TP-compounds with 7 doses and last raw as vehicle control (PBS). Shift in the EC_50_ were calculated. For IC_50_ determination of TP1 compound, we use EC_20_ of AngII the dose-response of TP1 in the drug plate. Fluorescence was recorded using a FlexStation 3 microplate reader (Molecular Devices) controlled by SoftMax® Pro software. Basal fluorescence was measured first recording 11 readings taken 5 s apart and instrument settings such as LED intensity and gain were adjusted to achieve the desired range (typical baseline readings at 10-20 RFU). AngII was then added at indicated concentrations using an automated liquid dispenser to minimize well-to-well variation. The fluorescence changes were recorded starting immediately and continued for 120 s until the signal returned to baseline to ensure the capture of peak response. The experiment utilized a non-cumulative dose-response protocol: Each Ang II concentration was applied to a separate triplicate set of wells to prevent receptor desensitization and AT1R tachyphylaxis. Intracellular calcium changes were quantified as ΔF/F (Fmax - Fmin) and reported as calcium response (RFUs) as authenticated in previous studies^17,63^. Assay variability was evaluated using response ΔF/F (Fmax - Fmin) in triplicate wells/dose of AngII across three independent days, and the average ± standard deviation was calculated to assess assay robustness. The EC₅₀/IC₅₀ values were derived from sigmoidal concentration-response curves fitted in GraphPad Prism 10 using data averaged from three independent experiments, each performed in triplicate.

### Cell Migration Assay

Mouse vascular aortic smooth muscle, MOVAS-AT1R cells cells were suspended 500,000 cells/ml in culture medium and 70μl were pipetted into each chamber of the cell culture insert (ibidi, Munich Germany). Cells were transferred to serum free medium 24 hours prior to removal of cell culture insert to ensure growth arrest. After removal of inserts, plates were washed with serum free media to remove cells which became detached prior to beginning treatments. Cells were treated with vehicle, [Sar1] AngII (1μM), [Sar1] AngII (1uM)+ Olm (1μM) or [Sar1] AngII (1μM) + TP1 (10μM). Live-cell images were acquired at 0, 8, and 24 hours. At 24 hours, cells were stained with DAPI to determine the number of cells that had migrated into the scratch area using 10X magnification (EVOS FL imaging system, ThermoFisher Scientific). The distance of migration and number of cells in the migrated zone in the images were analyzed using ImageJ software (NIH, Rockville, USA). Statistical analysis was performed using one-way ANOVA with Tukey’s multiple-comparison test. Data was normalized to the untreated control, which was set at 100%.

### Saturation binding

Saturation binding assays were performed using ¹²⁵I-AngII at concentrations ranging from 0.125 to 12 nM, in the absence or presence of TP1 (20 μM). Binding reactions were carried out in triplicate in 96-well plates and incubated for 1 h at room temperature to reach equilibrium. Total binding at each ¹²⁵I-AngII concentration was determined, and equilibrium dissociation constant (K_d_) and maximum binding capacity (B_max_) values were derived by nonlinear regression curve fitting in GraphPad Prism 10, using a one-site specific binding model. Data are presented as mean ± SEM^17,42^.

### Construction of AT1R mutants and characterization

Complementary DNA (cDNA) encoding wild-type AT1R and its alanine mutants (H166A, F170A, H183A, E185A, Q187A, N188A, and T190A), each with an N-terminal HA-tag, was synthesized and cloned into the expression vector pcDNA3.1 by GenScript (USA). HEK293T cells were grown in Dulbecco’s Modified Eagle’s Medium (DMEM; Invitrogen) supplemented with 10% fetal bovine serum (FBS; Thermo Fisher Scientific). Cells were seeded onto poly-D-lysine-treated culture plates at a density of 3 × 10⁶ cells per 10-cm plate. Following overnight culture, cells were transiently transfected with wild-type or mutant AT1R DNA using FuGENE 6 transfection reagent (Roche Applied Science) according to the manufacturer’s instructions. At 60-70 h post-transfection, cells were harvested and total membranes were prepared for radioligand binding analysis as described above^10,63–65^.

### Single-Point Radioligand Binding in AT1R Alanine Mutants and AT2R Cross-Reactivity Assessment

Membranes prepared from HEK293T cells transiently expressing wild-type or mutant AT1R (H166A, F170A, H183A, E185A, Q187A, or T190A) were used to assess the effect of TP1 on ¹²⁵I-AngII binding. Membranes were incubated with ¹²⁵I-AngII at a single concentration of 12 nM, the concentration identified from the wild-type saturation binding analysis as producing the largest and most statistically significant difference in total binding between AngII alone and AngII plus TP1, in the absence or presence of TP1 (20 μM), for 1 h at room temperature. Reactions were performed in triplicate for each construct, and the effect of TP1 on ¹²⁵I-AngII binding was compared between WT and each mutant by unpaired t-test^17^.

To assess TP1 selectivity for AT1R over AT2R, the same single-point binding protocol was performed using membranes from cells expressing AT2R. ¹²⁵I-AngII (12 nM) binding was assessed in the absence or presence of TP1 (20 μM), the AT1R-selective antagonist olmesartan, the AT2R-selective antagonist PD123,319, or unlabeled AngII, with olmesartan and PD123,319 included as reciprocal specificity controls in both the AT1R and AT2R binding systems. All conditions were performed in technical quadruplicate (n = 4 per condition), and the entire experiment was independently repeated three times, with a representative experiment shown. Statistical comparisons across conditions were performed by one-way ANOVA with post hoc multiple comparisons testing.

### Vascular Reactivity

Vascular contraction studies were conducted in accordance with National Institutes of Health guidelines for the care and use of laboratory animals and under protocols approved by the Institutional Animal Care and Use Committee. On the day of experimentation, male C57BL/6 mice were euthanized by isoflurane overdose followed by cervical dislocation. Iliac artery segments (1.5-2 mm) were excised, transferred to physiological saline solution (PSS: 130 mM NaCl, 4.7 mM KCl, 1.18 mM KH₂PO₄, 1.17 mM MgSO₄, 14.9 mM NaHCO₃, 5.5 mM dextrose, 26 μM CaNa₂ Versenate, 1.6 mM CaCl₂), and cleared of perivascular fat. Vessels were mounted on a multi-wire myograph system (Danish Myo Technologies, model 620N) using 40 μm stainless steel wires and incubated in warmed, aerated PSS (95% O₂/5% CO₂) for 60 minutes. Passive resting tension was normalized, and vessels were allowed to equilibrate for 20 minutes prior to functional testing.

Vessel integrity was assessed by two complementary approaches: (a) voltage-dependent contraction, induced by receptor-independent depolarization of vascular smooth muscle via elevated extracellular K⁺. PSS was replaced with a 60 mM K⁺ isotonic depolarizing buffer (KPSS: 74.7 mM NaCl, 60 mM KCl, 1.18 mM KH₂PO₄, 1.17 mM MgSO₄, 14.9 mM NaHCO₃, 5.5 mM dextrose, 26 μM CaNa₂ Versenate, 1.6 mM CaCl₂); following the contraction plateau, KPSS was washed out and replaced with PSS; and (b) receptor-dependent contraction and relaxation, assessed by α1-adrenergic and muscarinic receptor stimulation using phenylephrine and methacholine, respectively. Phenylephrine (200 nM) was added to the PSS-containing chamber, and the resulting contraction was relaxed by subsequent addition of methacholine (1 μM). Only vessels that passed both integrity tests were used to assess AngII-dependent contraction. Contractile responses were recorded using LabChart 8 data acquisition software (AD Instruments, Colorado Springs, CO)^17,42,62^.

### AngII-Dependent Contraction

Vessels that passed integrity testing were pre-incubated with either vehicle or TP1 (20 μM) for 5 minutes prior to stimulation with AngII (1 nM). Contractile responses were normalized to the KPSS-induced maximal contraction obtained for each vessel during integrity testing and expressed as percent contraction (%KPSS). AngII-induced contraction was compared between the vehicle (AngII-alone) and TP1 (AngII+TP1) groups, using independent vessels for each condition (n = 8 vessels from N = 4 animals per group), by unpaired t-test^17,42,62^.

### Statistical Considerations

Sample sizes (number of animals) were not predetermined by a formal statistical power calculation, and animals were used randomly. To prevent potential vessel damage during mounting, a defined exclusion criterion was applied: vessels lacking an adequate AngII-induced contraction, based on assessment of KPSS, phenylephrine, and methacholine responses, were excluded from analysis. Statistical analyses were performed after assessing normality of data distribution and homogeneity of variance for each dataset.

### Animal Care and Use

Male C57BL/6J were purchased from Jackson Laboratory (Bar Harbor, ME). Mice were cared for in accordance with the Guide for the Care and Use of Laboratory Animals in the Biological Resources Unit. All mouse experimental protocols described were reviewed and approved specifically for this project by the Institutional Animal Care and Use Committee at the Cleveland Clinic. Mice are cared for and monitored for signs of distress by trained veterinarians and technical staff who are on duty 7 days of the week.

### Acute Ligand Administration in C57BL6/J Mice

Ten-week-old male C57BL6/J mice were used for radiotelemetry transducer implantation. Anesthesia was induced and maintained by intraperitoneal injection of ketamine (100 mg/kg) and xylazine (10 mg/kg). Following isolation of the left common carotid artery, the transducer catheter was introduced into the carotid artery and advanced until its tip lay just within the thoracic aorta; the transmitter body (PA-C10, DSI) was positioned subcutaneously along the right flank. Postsurgical analgesia consisted of subcutaneous injection of flunixin (2.5 mg/kg). Mice were allowed to recover for 10 days prior to acute ligand administration.

Male mice implanted with telemetry transducers (n = 3) were tested for acute blood pressure responses to a 50 μL intraperitoneal bolus injection of vehicle, AngII (0.2 mg/kg), AngII (0.2 mg/kg) plus olmesartan (0.4 mg/kg), or AngII (0.2 mg/kg) plus TP1 (0.4 mg/kg). Systolic blood pressure (SBP), diastolic blood pressure (DBP), and heart rate (HR) were recorded for 30 minutes prior to injection to establish baseline values and for 45 minutes following injection to capture the induced blood pressure response, for a total recording period of 75 minutes per session. Because the magnitude and timing of the peak blood pressure response can be influenced by variability in absorption and hepatic metabolism, each treatment condition was independently repeated three times per animal, with a minimum 48-hour washout period between sessions to allow clearance of injected ligands, yielding a total of nine SBP and DBP recordings per treatment group (3 animals × 3 repeated sessions). This repeated-testing approach was used to assess the reproducibility and robustness of the acute response within and across animals, and to confirm AT1R specificity of the observed responses across independent trials of AngII, AngII plus olmesartan, and AngII plus TP1.

### Telemetry Data Analysis

Blood pressure was recorded at 2-minute intervals using Ponemah software (DSI, New Brighton, MN). Correct radiotelemetry sensor placement was confirmed by a pulse pressure greater than 15 mmHg and a systolic blood pressure between 80 and 200 mmHg, indicating a non-occluded catheter; recordings not meeting these criteria were excluded from analysis. SBP and DBP were recorded for 45 minutes post-injection in conscious, freely moving mice without anesthesia, with access to food and water restricted during the recording period. Blood pressure typically reached a maximum approximately 10 minutes post-injection and gradually declined toward baseline thereafter.

For each treatment group, individual recordings (3 animals per group, with acute responses independently assessed across repeated sessions per animal, separated by a minimum 48-hour washout) were averaged at each time point to generate a group mean time course. A subset of recordings was excluded from analysis due to technical issues (e.g., signal artifact or catheter occlusion), yielding a final count of 9 recordings for the vehicle, AngII, AngII+olmesartan, and AngII+TP1 groups, respectively. The decline in blood pressure was quantified from the group mean time course by averaging SBP andDBP over the final 18 minutes of the recording period (nine 2-minute time points) and comparing this value across treatment groups relative to the AngII-alone group, using one-way ANOVA followed by Dunnett’s multiple comparisons test (GraphPad Prism 10).

### Human Serum Sample Use

De-identified serum samples collected from consented participants under protocols approved by the Cleveland Clinic IRB were utilized for this study. Serum samples from patients with DCM were collected under IRB 06-805 and serum samples from preeclampsia patients were collected under IRB FA0901001. The serum samples were frozen in aliquots at -80°C after collection and thawed prior to use in the cell-ELISA (cELISA) experiments. Associated deidentified personal and clinical information for the samples was provided by our collaborator, Dr. W. H. Wilson Tang.

### Cell-Based ELISA (cELISA)

HA-AT1R-expressing cells were seeded at a density of 50,000 cells/well in poly-L-lysine-coated, clear-bottom 96-well plates and incubated overnight. Plates were washed with Wash Buffer 1 (WB1; 1% BSA in HBSS) and cells were fixed with 4% paraformaldehyde for 15 minutes at room temperature, followed by two additional washes with WB1. Following incubation with test compound or competing peptide (described for each assay below), primary antibody, patient serum or monoclonal antibody, as applicable, was added at a 1:1000 dilution and incubated for 1 hour at room temperature. Plates were washed with WB1, and HRP-conjugated anti-human secondary antibody (for patient serum samples) or HRP-conjugated anti-rat secondary antibody (for the monoclonal antibody) was added and incubated for 1 hour at room temperature. Plates were washed with Wash Buffer 2 (WB2; 0.5% BSA in HBSS), and 200 μL of OPD substrate solution (o-phenylenediamine dihydrochloride in phosphate-citrate buffer with sodium perborate) was added to each well; color development was allowed to proceed for 15 minutes and stopped with 3N HCl. Absorbance was measured at 492 nm using a FlexStation 3 Multi-Mode Microplate Reader (Molecular Devices)^17^.

### Screening of AT1R Autoantibodies in PE and DCM Patient Serum

Serum samples from patients with preeclampsia (PE; n = 84) and dilated cardiomyopathy (DCM; n = 43) were screened for AT1R autoantibody (AT1R-AAb) reactivity using the general cell-based ELISA protocol described above, with serum diluted 1:50 in WB1 in place of TP1 or competing peptide pre-incubation. The three highest-titer samples from each disease cohort were selected for further characterization.

### ECL2 Peptide Competition Assay

To confirm that the identified high-titer AT1R-AAb were directed against the ECL2 of AT1R, the three highest-titer serum samples from each disease cohort were pre-incubated with a competing AT1R ECL2 peptide prior to serum addition, following the general protocol above. Pre-incubation with the ECL2 peptide significantly reduced antibody binding relative to serum alone across all six samples tested, confirming that these autoantibodies are directed against the ECL2 epitope of AT1R.

### TP1 Competition Assay

To assess whether TP1 disrupts AT1R-AAb binding, the six high-titer, ECL2-confirmed serum samples (three PE, three DCM) were used in a dose-response competition format. Cells were pre-incubated for 1 hour at room temperature with TP1 across an eight-point, 2.5-fold serial dilution series (10, 4, 1.6, 0.64, 0.256, 0.1024, 0.04096, and 0.016384 μM) prior to addition of patient serum (1:1000), following the general protocol above, with detection by HRP-conjugated anti-human secondary antibody. For the monoclonal antibody directed against the AT1R ECL2 epitope, the same competition format was used with TP1 tested across a broader concentration range, with detection by HRP-conjugated anti-rat secondary antibody. Percent inhibition of antibody binding by TP1 was calculated relative to the no-TP1 control for each sample. Data were normalized and analyzed using the sigmoidal dose-response (variable slope) function in GraphPad Prism 10 to derive IC_50_ values.

### Cytotoxicity and Cell Viability Assays

Cell viability and cytotoxicity were assessed in HEK293T cells stably expressing AT1R (HEK293T-AT1R) following 72 hours of treatment with vehicle, AngII (20 μM; Bachem, Torrance, CA), olmesartan (20 μM; Daiichi Sankyo Chemical Pharma), or TP1 (0.2, 2, or 20 μM). Cell viability was measured using the CellTiter-Blue Cell Viability Assay (Promega, catalog #G8080), and cytotoxicity was assessed in parallel using the CellTox Green Cytotoxicity Assay (Promega, catalog #G8741), according to the manufacturer’s instructions. Lysis buffer, provided as part of the CellTox Green Cytotoxicity Assay kit, served as a positive control for the cytotoxicity assay. Each experiment was independently performed three times on separate days, with eight technical replicates per condition for the viability assay and four technical replicates per condition for the cytotoxicity assay. Data were analyzed by one-way ANOVA with Dunnett’s multiple comparisons test, comparing each treatment group to vehicle, with p < 0.05 considered statistically significant.

### Pharmacokinetic and Physicochemical Characterization of TP1

Kinetic aqueous solubility of TP1 was determined in PBS (pH 7.4) by DMSO-dilution shake-flask assay with LC-MS/MS quantification (100 μM nominal concentration, n = 3). Bidirectional Caco-2 cell permeability was assessed using confluent monolayers (passage 40) with atenolol, propranolol, and digoxin as low-permeability, high-permeability, and P-glycoprotein (P-gp) substrate reference compounds, respectively (10 μM, 120 min, 37°C, n = 2 per direction); apparent permeability (Papp) and efflux ratio were calculated as described in SI Appendix, Methods. Metabolic stability was assessed in rat and human liver microsomes (0.5 mg/mL protein, 1 μM compound, NADPH-supplemented, 37°C, 0-45 min, N = 2), with ketanserin as reference compound; in vitro half-life, intrinsic clearance (CL_int_), and hepatic extraction ratio (E*_h_*) were derived from first-order elimination kinetics using the well-stirred liver model. Plasma protein binding was determined by equilibrium dialysis in human plasma (5 μM, 37°C, 5 h, n = 2), with warfarin as a high-protein-binding reference compound. Single-dose pharmacokinetics were evaluated in male Sprague-Dawley rats (n = 3/group) following intravenous (2 mg/kg) or oral (10 mg/kg) administration in aqueous vehicle (5% DMSO, 10% Solutol, 85% saline); plasma concentrations were determined by LC-MS/MS, and PK parameters were derived by noncompartmental analysis (Phoenix WinNonlin 7.0), including absolute oral bioavailability (F). Full assay protocols, calculation formulas, and classification thresholds are provided in **SI, Methods**.

## Data availability

All data generated or analyzed during this study are included in this published article and the supplementary information provided online. Further queries may be addressed to the corresponding author Dr. Sadashiva Karnik.

## Acknowledgements

The authors acknowledge Cleveland Clinic Innovations and Core services including the Animal Core Facility, the Histopathology and Imaging Core; Lousie Lantier and the Vanderbilt University Mouse Metabolic Phenotyping Center (RRID: SCR_021939) for assistance with the telemetry blood pressure study; This work was supported by National Institutes of Health (NIH) RO1 grants, HL142091 and HL132351 and Lerner Research Institute’ Catalyst Award to S.S. K, and Center of Excellence grant to WHWT and SSK. to SSK. The Vanderbilt Mouse Metabolic Phenotyping Center is supported by NIH grants, DK135073, DK020593.

## Author contributions

KDS, TRP, SSK carried out compound design, computational docking and mutagenesis, Experimental validation study design and execution and data analyses were done by TRH, RD, LL, TR and KDS and were supervised by SSK. APM and WHWT provided control and patient derived autoantibodies for this study. KDS, and SSK wrote the manuscript, with critical input from all co-authors, who have read, revised and approved the manuscript.

## Disclosures

The authors declare no conflict of interests.

## Additional information

Online Supplementary Information is available.

## Notes

### Competing Interest Statement

The authors have declared no competing interest.

