## Supplemental Material for "AT1R-Selective Negative Allosteric Modulator Targeting the Epitope for Pathogenic Autoantibody Activation"

**HBKR Department, NB5-76**

**Cleveland Clinic Research,**

**9500 Euclid Avenue**

**Cleveland, OH 44195**

**U.S.A.**

**216-444-1269 (O)**

### **Supplementary Results**

**TP1 Exhibits Adequate Kinetic Aqueous Solubility.** TP1 showed a mean kinetic solubility of $82.8 \pm 18.7 \mu\text{M}$ ($n = 3$; individual values 64.6, 81.9, 102 $\mu\text{M}$) in PBS (pH 7.4), corresponding to ~83% of the nominal 100 $\mu\text{M}$ test concentration remaining in solution (Table S1). This favorable solubility profile indicates that aqueous solubility is unlikely to be a limiting factor for TP1's in vitro potency or in vivo formulation, consistent with successful formulation of TP1 in aqueous vehicle for PK and pharmacology studies.

**TP1 Shows Low Passive Permeability and Evidence of P-gp-Mediated Efflux. Reference compounds validated the Caco-2 assay:** atenolol showed low permeability ($P_{\text{app A} \rightarrow \text{B}} = 1.15 \times 10^{-6} \text{ cm/s}$), propranolol showed high permeability ($19.98 \times 10^{-6} \text{ cm/s}$), and digoxin showed low passive permeability ($0.67 \times 10^{-6} \text{ cm/s}$) with a high efflux ratio (23.9), consistent with its established P-gp substrate status (Table S2). TP1 showed low A→B permeability ($1.43 \times 10^{-6} \text{ cm/s}$) and a high efflux ratio (12.9), with acceptable recovery in both directions (93.3–101.4%), classifying it as having low passive permeability and potential P-gp substrate behavior, factors likely contributing to its limited oral bioavailability.

**TP1 Exhibits Moderate-to-High Metabolic Clearance in Liver Microsomes.** In rat liver microsomes, TP1 was metabolized more rapidly than ketanserin (12.3% vs. 28.5% remaining at 45 min), corresponding to $t_{1/2}$ of 15.0 vs. 24.6 min and Eh of 66.1% vs. 54.4% (Table S3, Fig. S1). In human microsomes, TP1 similarly cleared faster than ketanserin (25.9% vs. 34.8% remaining at 45 min; $t_{1/2}$ 25.9 vs. 34.8 min; Eh 74.9% vs. 69.0%). Ketanserin half-lives matched historical reference values in both species, confirming assay validity. No-cofactor controls confirmed that TP1 depletion in rat microsomes was predominantly NADPH-dependent (97.4% remaining without NADPH); a partial NADPH-independent component was noted in human microsomes (44.6% remaining without NADPH at 45 min), a caveat for interpreting the human CL<sub>int</sub>/Eh values.

**TP1 Shows Moderate Plasma Protein Binding in Human Plasma.** TP1 was moderately protein-bound in human plasma (88.6% bound; $F_u = 11.4\%$; Table S4), while warfarin showed high binding (99.4%), consistent with historical values, confirming assay validity. TP1's moderate protein binding supports adequate free-drug availability at pharmacologically active concentrations.

**TP1 Exhibits High Clearance and Low Oral Bioavailability in Rats.** Following IV dosing, TP1 showed a short half-life ($T_{1/2} = 0.47 \pm 0.27 \text{ h}$), high clearance ($\text{Cl} = 460.2 \pm 183.6 \text{ mL/h/kg}$), and large volume of distribution ($V_{\text{ss}} = 6671 \pm 3498 \text{ mL/kg}$) (Table S5). Following oral dosing, TP1 showed low absolute bioavailability ($F = 15.3 \pm 1.9\%$), a longer apparent terminal half-life ($T_{1/2} = 1.19 \pm 0.14 \text{ h}$, likely reflecting flip-flop kinetics), and a secondary rise in plasma concentration between 2–4 h post-dose suggestive of possible double-peak absorption behavior. These findings are consistent with the moderate-to-high hepatic extraction and low Caco-2 permeability/high efflux ratio observed in vitro and collectively explain TP1's limited oral exposure.

**Table S1. Kinetic aqueous solubility of TP1 in PBS (pH 7.4).**

| Compound | Nominal Conc. (μM) | Rep 1 (μM) | Rep 2 (μM) | Rep 3 (μM) | Mean ± SD (μM) | % of Nominal |
| --- | --- | --- | --- | --- | --- | --- |
| TP1 | 100 | 64.6 | 81.9 | 102 | 82.8 ± 18.7 | 82.8% |

*Kinetic solubility is determined by shake-flask method with LC-MS/MS quantification following 1 h equilibration in PBS (pH 7.4) at room temperature. n = 3 independent replicate incubations.*

**Table S2. Caco-2 bidirectional permeability of TP1 and reference compounds.**

| Compound | Papp A→B (×10<sup>-6</sup> cm/s) | Papp B→A (×10<sup>-6</sup> cm/s) | Efflux Ratio | Recovery A→B (%) | Recovery B→A (%) | Classification | P-gp Substrate |
| --- | --- | --- | --- | --- | --- | --- | --- |
| Atenolol | 1.15 | NA | NA | 103.7 | NA | Low | No |
| Propranolol | 19.98 | NA | NA | 74.3 | NA | High | No |
| Digoxin | 0.67 | 16.09 | 23.9 | 100.7 | 101.4 | Low | Yes (established) |
| TP1 | 1.43 | 18.53 | 12.9 | 93.3 | 101.0 | Low | Potential |

*Papp, apparent permeability coefficient; test concentration, 10 μM; incubation, 120 min at 37°C; n = 2 replicate wells per direction. Permeability classification: low, Papp(A→B) < 2 × 10<sup>-6</sup> cm/s; moderate, 2–10 × 10<sup>-6</sup> cm/s; high, >10 × 10<sup>-6</sup> cm/s.*

**Table S3. Microsomal stability parameters for TP1 and ketanserin in rat and human liver microsomes.**

| Compound | Species | % Rem. (0 min) | % Rem. (5 min) | % Rem. (15 min) | % Rem. (30 min) | % Rem. (45 min) | t<sub>1/2</sub> (min) | CL<sub>int</sub> (mL/min/kg) | CL<sub>h</sub> (mL/min/kg) | E<sub>h</sub> (%) |
| --- | --- | --- | --- | --- | --- | --- | --- | --- | --- | --- |
| Ketanserin | Rat | 100 | 88.0 | 67.3 | 42.6 | 28.5 | 24.6 | 46.3 | 46.3 | 54.4 |
| TP1 | Rat | 100 | 65.7 | 33.8 | 18.0 | 12.3 | 15.0 | 165.8 | 56.2 | 66.1 |
| Ketanserin | Human | 100 | 83.1 | 68.5 | 49.6 | 40.3 | 34.8 | 14.3 | 14.3 | 69.0 |
| TP1 | Human | 100 | 79.2 | 50.9 | 38.4 | 29.1 | 25.9 | 61.9 | 15.5 | 74.9 |

Data represent mean values from N = 2 replicate incubations per time point. Test concentration, 1 μM; microsomal protein concentration, 0.5 mg/mL. t<sub>1/2</sub>, in vitro half-life; CL<sub>int</sub>, intrinsic clearance; CL<sub>h</sub>, predicted hepatic clearance (well-stirred model); E<sub>h</sub>, predicted hepatic extraction ratio.

**Table S4. Plasma protein binding of TP1 and warfarin in human plasma.**

| Test Article | Bound Fraction (%) | F<sub>u</sub> (%) | Category | Recovery (%) |
| --- | --- | --- | --- | --- |
| Warfarin | 99.4 | 0.6 | High | 94.3 |
| TP1 | 88.6 | 11.4 | Moderate | 83.4 |

Test concentration, 5 μM; equilibrium dialysis, 5 h at 37°C; n = 2 replicate incubations. Category thresholds: high, >90% bound; moderate, 50-90% bound; low, <50% bound.

**Table S5. Pharmacokinetic parameters of TP1 following single IV (2 mg/kg) and PO (10 mg/kg) administration in rats.**

| Parameter | IV, Mean ± SD | PO, Mean ± SD |
| --- | --- | --- |
| $T_{1/2}$ (h) | 0.47 ± 0.27 | 1.19 ± 0.14 |
| $T_{max}$ (h) | 0.083 (nominal) | 2.75 ± 2.17 |
| $C_{max}$ (ng/mL) | 253.9 ± 125.4 | 23.1 ± 5.5 |
| AUC<sub>0-t</sub> (h·ng/mL) | 81.5 ± 35.9 | 62.5 ± 7.8 |
| AUC<sub>0-∞</sub> (h·ng/mL) | 81.7 ± 35.9 | 65.5 ± 8.6 |
| Cl (mL/h/kg) | 460.2 ± 183.6 | - |
| $V_{ss}$ (mL/kg) | 6671 ± 3498 | - |
| F (%) | - | 15.3 ± 1.9 |

*n* = 3 animals per group (male Sprague-Dawley rats). $T_{max}$ reported as nominal first sampling time (IV) or mean ± SD (PO). PK parameters derived by noncompartmental analysis in Phoenix WinNonlin 7.0. Cl and $V_{ss}$ reported for IV only; F reported for PO only, per standard noncompartmental PK convention.

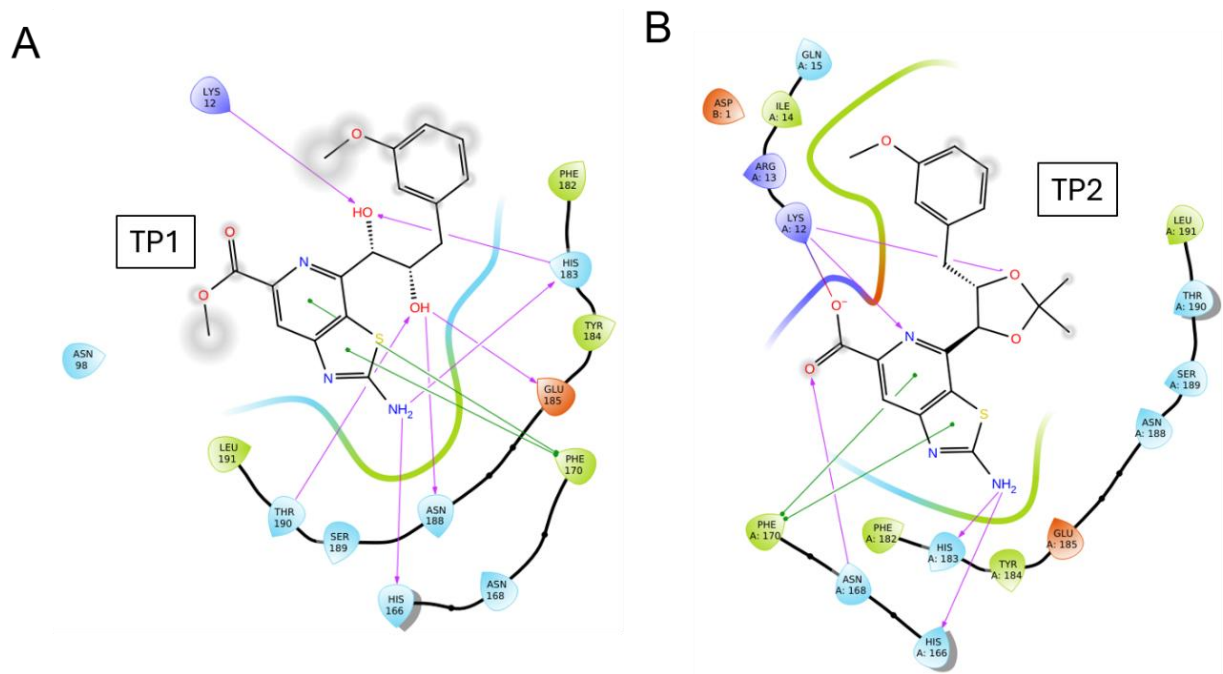

**Figure S1. Two-dimensional ligand interaction diagrams and docking scores for TP1 and TP2.** 2D interaction diagrams generated from Glide XP docking of TP1 (left) and TP2 (right) at the predicted allosteric site of the AT1R-AngII complex. Hydrogen bonds are shown as pink arrows/dashed lines; residues contributing to the binding pocket are labeled and colored by residue type (polar, hydrophobic, acidic, basic). TP1 engages the pocket through direct hydrogen bonding contacts mediated by the free vicinal diol, achieving a Glide docking score of -9.51 kcal/mol. Acetonide protection of the diol in TP2 disrupts this hydrogen bonding network, resulting in a reduced docking score of -7.54 kcal/mol.

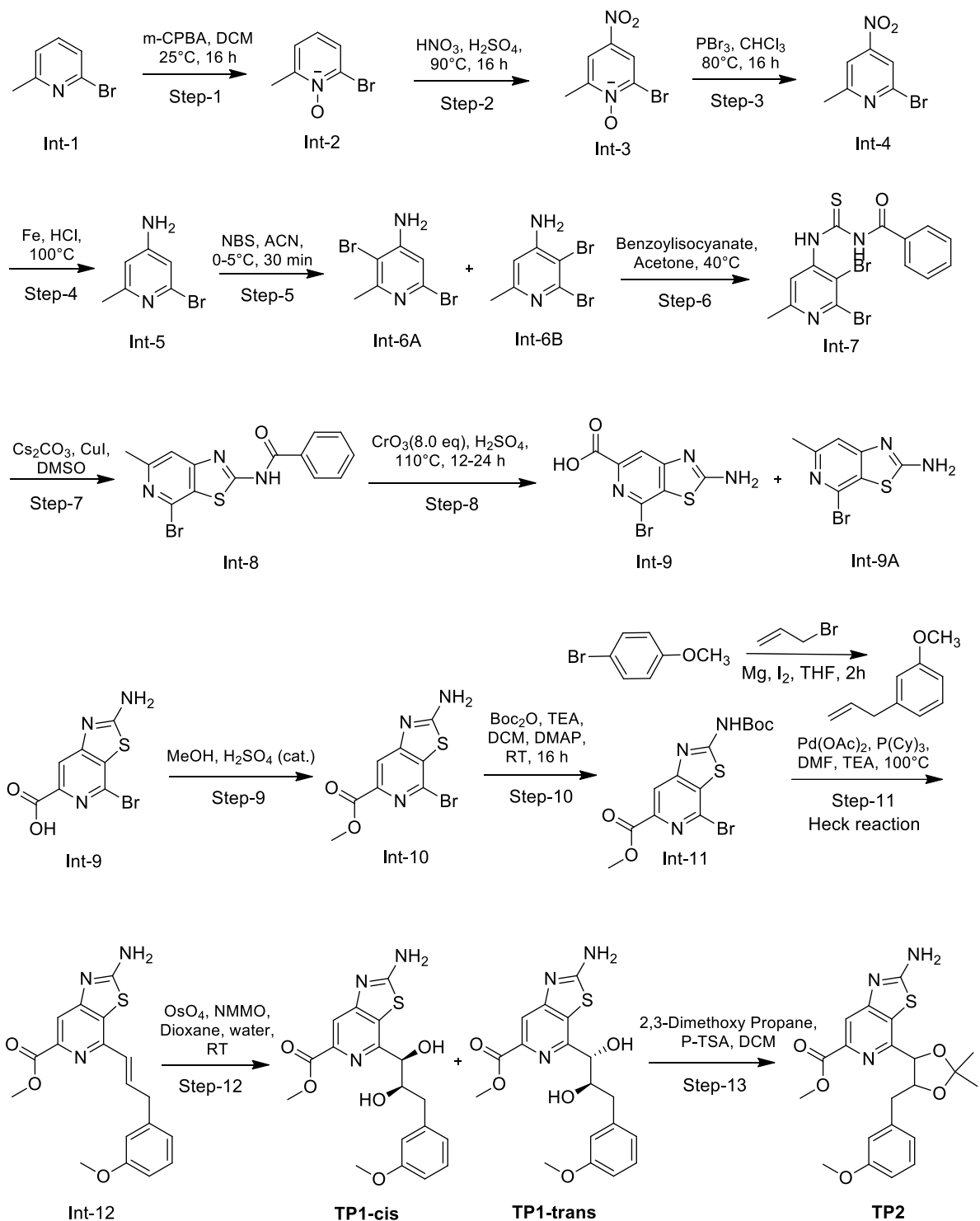

**Figure S2. Hypothesis-Driven Synthetic Route to TP1 and TP2.** Following structure-based design of TP1 as an optimized lead candidate built on a thiazolo[4,5-c]pyridine scaffold (**Fig. 1**), we developed a modular synthetic route to access this compound and its diol-protected analogue, enabling both physicochemical tuning and mechanistic evaluation of the key functional motifs identified through molecular docking.

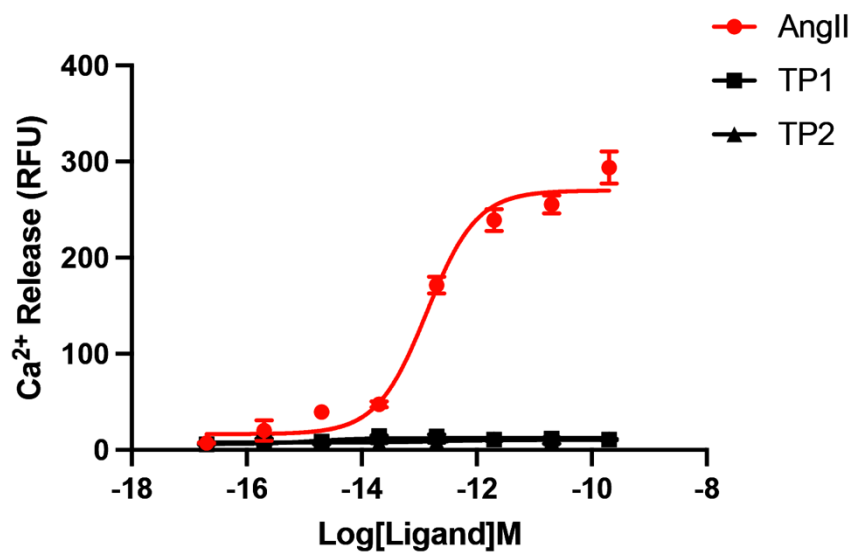

**Figure S3. TP1 and TP2 exhibit no intrinsic agonist activity at AT1R.** Concentration-response curves for intracellular $\text{Ca}^{2+}$ release (relative fluorescence units, RFU) in cells stimulated with AngII, TP1, or TP2 alone, in the absence of AngII co-stimulation. AngII produced the expected concentration-dependent increase in $\text{Ca}^{2+}$ release. In contrast, neither TP1 nor TP2 elicited any measurable $\text{Ca}^{2+}$ response across the concentration range tested, confirming that both compounds lack intrinsic agonist activity at AT1R and act exclusively as negative allosteric modulators of AngII-induced signaling rather than through direct receptor activation.

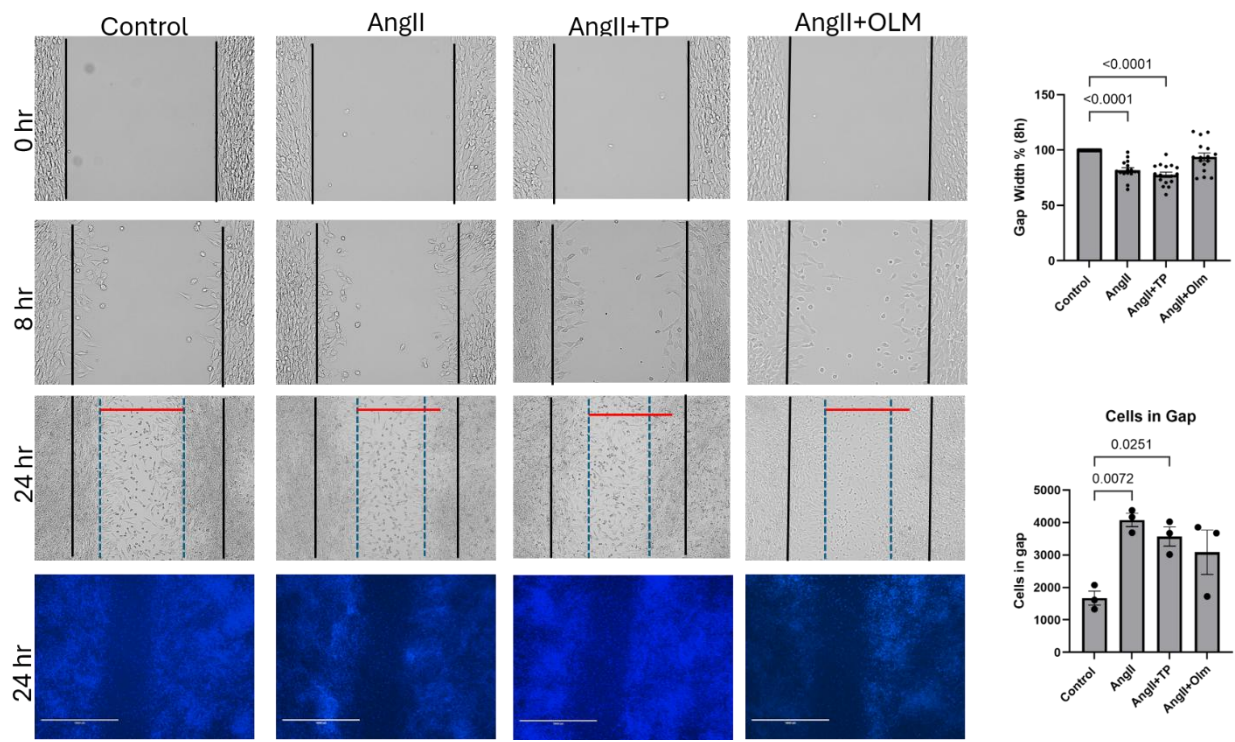

**Figure S4. Assessment of TP1 effect on vascular smooth muscle cell migration (proliferation) and scratch wound closure.** Mouse Vascular Arterial Smooth Muscle (MOVAS-AT1R) cells serum starved for 24hours, were treated with vehicle, [Sar1] AngII (1μM), [Sar1] AngII (1μM)+ Olm (1μM ) or [Sar1] AngII (1μM) + TP (10μM) and monitored for migration over 24 hours under serum free conditions. Images were acquired at 0, 8 and 24h, then stained with DAPI. After 24h, the number of cells in the migrated zone were analyzed using ImageJ. Statistics were performed by One Way ANOVA with Tukey's multiple comparison test and normalized to Control =100%. Change in distance traveled. Statistics were performed by One Way ANOVA with Dunnett's multiple comparison test. N=3 with 3 images per treatment per time point. Scale bar, 400 μm.

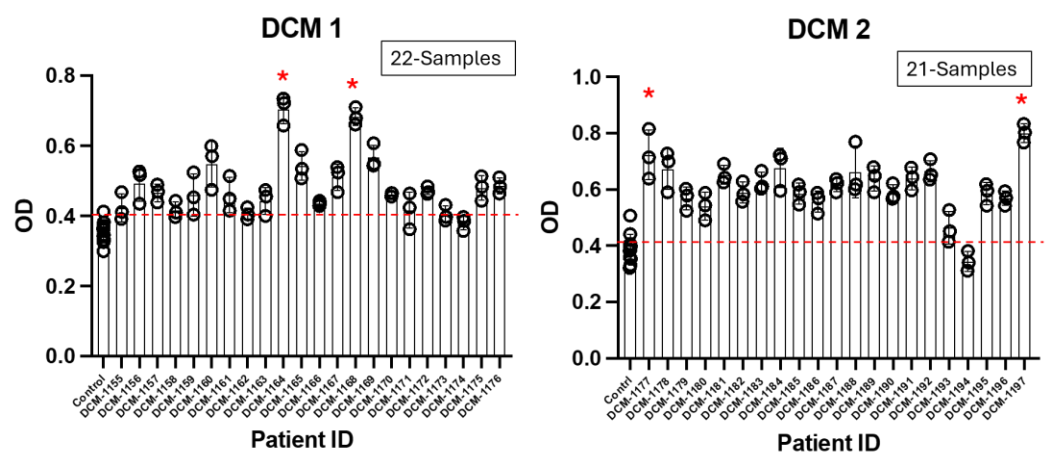

**Figure S5A. Screening identifies AT1R autoantibody positive serum samples in the DCM patient cohort.** Serum samples from patients with dilated cardiomyopathy (DCM, $n = 43$ total, divided into two screening cohorts: DCM 1, 22 samples; DCM 2, 21 samples) were screened for AT1R autoantibody (AT1R-AA) reactivity using a cell-based ELISA on AT1R-expressing cells fixed with 4% formaldehyde, measured as optical density (OD). Control represents pooled serum from healthy individuals. Data are presented as mean ± SEM from $n = 3$ technical replicates per sample.

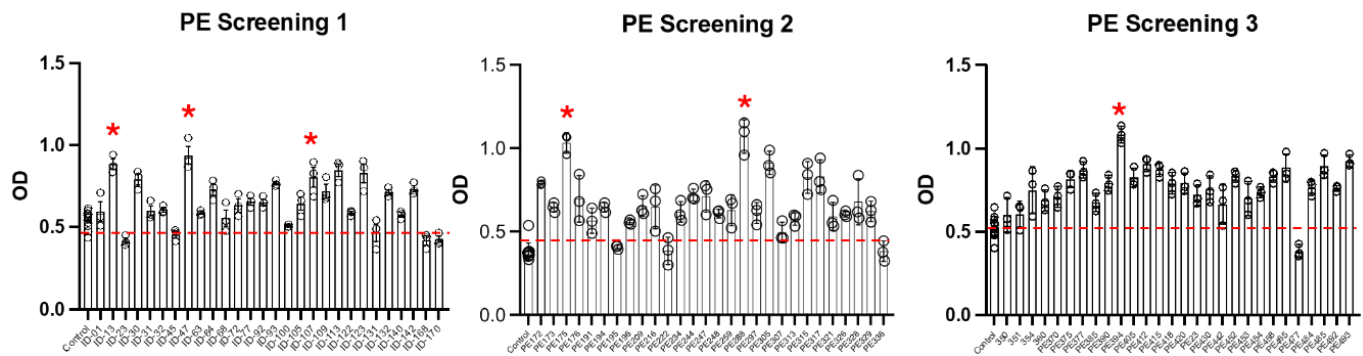

**Figure S5B. Screening identifies AT1R autoantibody reactivity in the PE patient cohort.** Serum samples from patients with preeclampsia (PE, $n = 84$ total, divided into three screening cohorts: PE Screening 1-3) were screened for AT1R autoantibody (AT1R-AA) reactivity using a cell-based ELISA on AT1R-expressing cells fixed with 4% formaldehyde, measured as optical density (OD). Control represents pooled serum from healthy individuals. Data are presented as mean ± SEM from $n = 3$ technical replicates per sample.

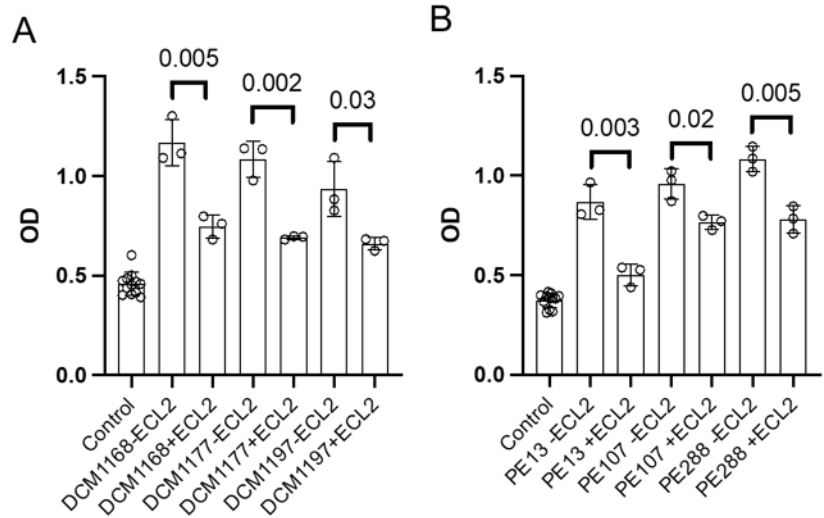

**Figure S6. AT1R autoantibodies from DCM and PE patients specifically target the ECL2 epitope of AT1R.** Cell-based ELISA showing binding of high-titer AT1R autoantibody (AT1R-AA) positive serum samples, from DCM patients (DCM1168, DCM1177, DCM1197; left) and PE patients (PE13, PE107, PE288; right), to AT1R-expressing cells in the absence or presence of competing AT1R extracellular loop peptides corresponding to ECL1, ECL2, or ECL3. ECL1 and ECL2 peptides did not significantly affect AT1R-AA binding (data not shown). Whereas pre-incubation with the ECL2 peptide significantly reduced serum antibody binding across all six samples tested (DCM1168, $p = 0.005$; DCM1177, $p = 0.002$; DCM1197, $p = 0.03$; PE13, $p = 0.003$; PE107, $p = 0.02$; PE288, $p = 0.005$; unpaired t-test). In contrast, pre-incubation with either the ECL1 or ECL3 peptide had no significant effect on antibody binding for any sample tested, confirming that these autoantibodies are specifically directed against the ECL2 epitope of AT1R rather than other extracellular loops. Control represents pooled serum from healthy individuals. Data are presented as mean ± SEM from $n = 3$ technical replicates per sample.

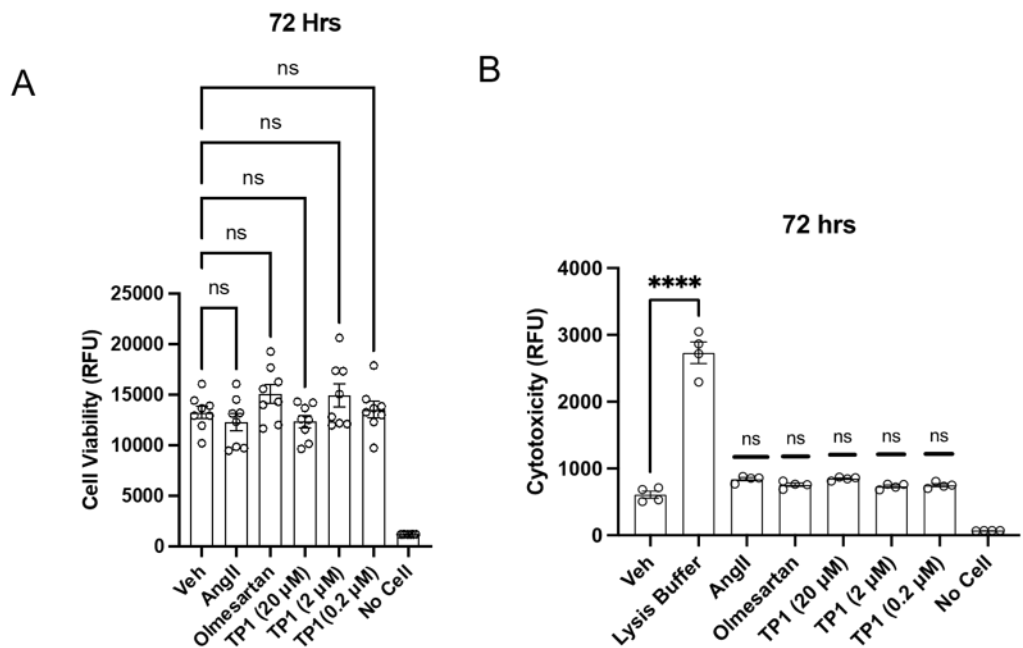

**Figure S7. TP1 does not impair cell viability or induce cytotoxicity in AT1R expressing HEK293T cells at 72 hours. (Left)** Cell viability, measured by CellTiter-Blue assay, in cells treated with vehicle, AngII (20 μM), olmesartan (20 μM), or TP1 (20, 2, or 0.2 μM) for 72 hours. **(Right)** Cytotoxicity, measured by CellTox Green assay, in the same treatment groups, with lysis buffer included as a positive control. No significant differences in viability or cytotoxicity were observed among vehicle, AngII, olmesartan, or any TP1 concentration. Lysis buffer produced a significant increase in cytotoxicity signal relative to vehicle (\*\*\*$p < 0.0001$). Data are shown as mean ± SEM from a representative experiment, independently repeated three times with consistent results. Statistical comparisons were performed by one-way ANOVA with Dunnett's multiple comparisons test relative to vehicle.

**A**

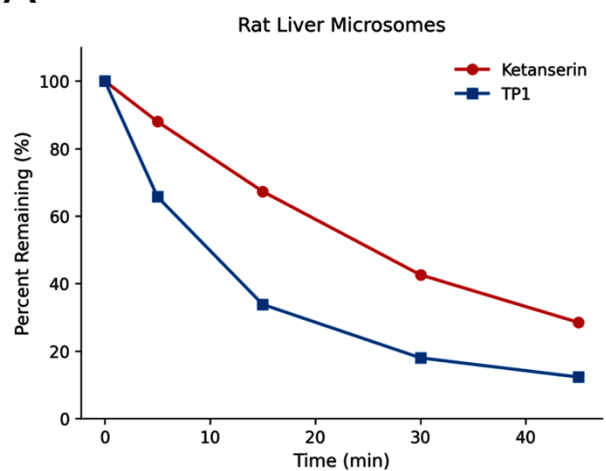

**B**

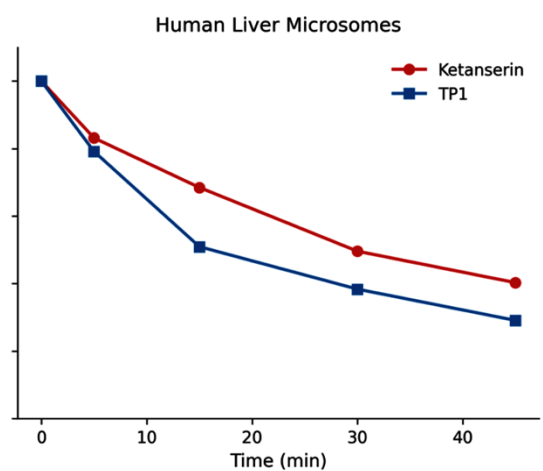

**Figure S8.** Percent parent compound remaining over a 45-minute incubation with NADPH-supplemented rat (left) or human (right) liver microsomes, for TP1 and ketanserin. n = 2 replicate incubations per time point.

### **Supplementary Methods:**

#### **Synthetic Creation of TP1 and TP2**

**Hypothesis-Driven Synthetic Route to TP1 and TP2.** Following structure-based design of TP1 as an optimized lead candidate built on a thiazolo[4,5-c]pyridine scaffold (Fig. 1), we developed a modular synthetic route to access this compound and its diol-protected analogue, enabling both physicochemical tuning and mechanistic evaluation of the key functional motifs identified through molecular docking.

**Phase I: Directed Functionalization of the Pyridine Core (Steps 1–4).** Synthesis commenced from commercially available 2-bromo-6-methylpyridine (Int-1), selected to provide a versatile halogen handle for regioselective diversification toward the target thiazolopyridine core. Oxidation of the pyridine nitrogen using *m*-chloroperbenzoic acid (*m*CPBA) afforded pyridine *N*-oxide (Int-2), which acted as a directing group to enable regioselective nitration at the C4 position (Int-3). Subsequent reductive deoxygenation restored the aromatic pyridine ring (Int-4), and chemoselective reduction of the nitro group yielded the key 4-amino-2-bromo-6-methylpyridine intermediate (Int-5), installing a nucleophilic primary amine handle essential for downstream heterocycle formation.

**Phase II: Assembly of the Thiazolo[4,5-c]pyridine Core (Steps 5–7).** Electrophilic bromination using *N*-bromosuccinimide (NBS) installed halogen substituents at activated positions, providing brominated intermediate species (Int-6) that contain orthogonal handles for annulation and late-stage cross-coupling. Reaction of the primary amine with benzoyl isothiocyanate generated a benzoylthiourea intermediate (Int-7). Subsequent copper-catalyzed intramolecular C–S bond formation and deprotection furnished the bicyclic 2-aminothiazolo[4,5-c]pyridine scaffold (Int-8), establishing the defining heterocyclic core designed to anchor the ligand within the AT1R allosteric domain.

**Phase III: Scaffold Diversification and Pharmacophore Installation (Steps 8–9A).** Selective oxidation of the benzylic C6-methyl substituent to a methyl ester (Int-9) introduced polar character and synthetic versatility aligned with the docking-predicted binding mode. Temporary protection of the exocyclic 2-amino group as a *tert*-butoxycarbonyl (Boc) carbamate (Int-9A) ensured chemoselectivity during subsequent cross-coupling. Installation of the 3-methoxystyryl side chain via palladium-catalyzed Heck cross-coupling with 3-methoxystyrene furnished Int-10, incorporating the 3-methoxyphenyl aromatic vector conserved from initial screening hits DCP1-3 and DCP1-16.

**Phase IV: Installation and Functional Validation of the Vicinal Diol Motif (Steps 10–12).** Upjohn catalytic osmium-mediated dihydroxylation of the styryl double bond in Int-10 yielded a mixture of vicinal diol diastereomers (Int-11-Fr-1 and Int-11-Fr-2, corresponding to TP1-cis and TP1-trans). This step installed the polar vicinal diol motif predicted by molecular docking to form crucial hydrogen bonds with allosteric pocket residues H183, E185, and Q187.

To directly probe this structural prediction, the vicinal diol was selectively masked via acid-catalyzed acetonide formation to afford the cyclic 1,3-dioxolane analogue (step-12 compound, designated TP2). Rather than a simple synthetic derivative, TP2 was deliberately synthesized as a structural probe to evaluate the contribution of diol-mediated hydrogen bonding to receptor engagement. Comparative biological evaluation of TP1 against TP2 confirmed that masking the free diol significantly reduces biological efficacy, corroborating the reduced Glide docking score predicted *in silico* for the acetonide-protected analogue.

**In Vitro Metabolic Stability in Liver Microsomes.** Metabolic stability of TP1 was assessed in rat and human liver microsomes (Xenotech), with ketanserin included as a reference compound of known metabolic clearance. Test and reference compounds were prepared as 500 $\mu\text{M}$ spiking solutions in DMSO from 10 mM stock solutions, then further diluted to a 1.5 $\mu\text{M}$ working solution in PBS (pH 7.4) containing 0.75 mg/mL liver microsomal protein. Reactions were initiated by combining 30 $\mu\text{L}$ of the 1.5 $\mu\text{M}$ compound/microsome solution with 15 $\mu\text{L}$ of NADPH stock solution (6 mM) to a final compound concentration of 1 $\mu\text{M}$ and a final microsomal protein concentration of 0.5 mg/mL and incubated at 37°C. Reactions were quenched at 0, 5, 15, 30, and 45 minutes by the addition of 150 $\mu\text{L}$ of acetonitrile:methanol (1:1) containing internal standard. A no-NADPH-cofactor (NC) control, in which PBS was substituted for NADPH, was included at the 0- and 45-minute time points to control for chemical (non-enzymatic) degradation. All incubations were performed in duplicate ($N = 2$). Following quenching, plates were shaken (600 rpm, 10 min) and centrifuged (6000 rpm, 15 min), and supernatants were diluted and analyzed by LC-MS/MS<sup>1</sup>.

Percent parent compound remaining at each time point was calculated relative to the 0-minute time point. The elimination rate constant ($K$) was determined from the slope of the linear regression of $\ln$ (percent remaining) versus incubation time, and in vitro half-life ($t_{1/2}$) was calculated as $t_{1/2} = 0.693/K$. Intrinsic clearance ($CL_{\text{int}}$) was calculated as $CL_{\text{int}} = (0.693/t_{1/2}) \times (1/\text{microsomal protein concentration}) \times \text{species-specific scaling factor (microsomal protein content per gram of liver} \times \text{liver weight per kilogram body weight)}$. Predicted hepatic clearance ($CL_{\text{h}}$) was calculated using the well-stirred liver model, $CL_{\text{h}} = CL_{\text{int}} \times Q_{\text{h}}/(Q_{\text{h}} + CL_{\text{int}})$, where $Q_{\text{h}}$ is species-specific hepatic blood flow (85 mL/min/kg for rat; 20.7 mL/min/kg for human). Hepatic extraction ratio ($E_{\text{h}}$) was calculated as $E_{\text{h}} (\%) = 100 \times CL_{\text{h}}/Q_{\text{h}}$. Compounds were classified as having low ($E_{\text{h}} \leq 30\%$), moderate ($30\% < E_{\text{h}} \leq 70\%$), or high ($E_{\text{h}} > 70\%$) predicted hepatic clearance<sup>2-4</sup>.

**Single-Dose Pharmacokinetics in Rats.** The pharmacokinetics of TP1 were evaluated in male Sprague-Dawley rats following single intravenous (IV) and oral (PO) administration. TP1 (free form, >98% purity) was formulated in 5% DMSO, 10% Solutol, and 85% saline for both routes. Animals ($n = 3$ per group) received TP1 by IV bolus at 2 mg/kg (dose volume 5 mL/kg) or by oral gavage at 10 mg/kg (dose volume 10 mL/kg); animals in the oral dosing group were fasted overnight (10-14 h) prior to dosing, with food access resumed 4 h post-dose. Serial blood samples (0.2 mL) were collected via jugular or other suitable vein into K2-EDTA tubes at 0.083, 0.25, 0.5, 1, 2, 4, 8, and 24 h post-dose for the IV group, and at 0.25, 0.5, 1, 2, 4, 6, 8, and 24 h post-dose for the PO group. Samples were centrifuged ($6800 \times g$, 6 min, 2-8°C) within 1 h of collection and plasma was stored at -80°C until analysis. Plasma TP1 concentrations were determined by LC-MS/MS, with quality control samples confirming intra-assay accuracy within 80-120% of nominal concentration for >66.7% of QC samples.

Pharmacokinetic parameters, including maximum plasma concentration ($C_{\text{max}}$), time to $C_{\text{max}}$ ($T_{\text{max}}$), terminal elimination half-life ($T_{1/2}$), area under the plasma concentration-time curve from time zero to the last measurable concentration ($AUC_{0-t}$) and extrapolated to infinity ($AUC_{0-\infty}$), clearance ($CL$), and steady-state volume of distribution ($V_{\text{ss}}$), were calculated by noncompartmental analysis using Phoenix WinNonlin (version 7.0; Pharsight, USA). Absolute oral bioavailability ($F$) was calculated as $F (\%) = [(AUC_{0-\infty}, \text{PO}/\text{Dose PO})/(AUC_{0-\infty}, \text{IV}/\text{Dose IV})] \times 100^{5-7}$.

**Plasma Protein Binding.** The extent of plasma protein binding of TP1 was determined by equilibrium dialysis in human plasma, with warfarin included as a reference compound of known, high plasma protein binding. Plasma was thawed at 37°C and centrifuged (12,000 rpm, 5 min) to remove clots prior to use. Test and reference compounds were spiked into plasma from a DMSO stock solution to a final test concentration of 5 μM (final DMSO concentration $\leq 1\%$, v/v). Spiked plasma (100 μL) was loaded into the donor chamber of a dialysis apparatus fitted with a pre-hydrated dialysis membrane, opposite 100 μL of blank dialysis buffer in the receiver chamber. An aliquot of the spiked plasma was reserved at the time of loading as the 0 h (T0) sample. Dialysis was performed at 37°C with shaking (60 rpm) for 5 h. At the end of the incubation, aliquots from the donor (plasma) and receiver (buffer) chambers were collected, mixed with an equal volume of the opposite blank matrix to normalize matrix effects, quenched with acetonitrile:methanol (1:1) containing internal standard, and analyzed by LC-MS/MS. All incubations were performed in duplicate (n = 2).

The fraction of compound bound to plasma protein was calculated as: % Bound = $100 \times ([\text{Donor}]5\text{h} - [\text{Receiver}]5\text{h})/[\text{Donor}]5\text{h}$, and the unbound fraction as $F_u (\%) = 100 - \% \text{ Bound}$. Assay recovery was calculated as % Recovery = $100 \times ([\text{Donor}]5\text{h} + [\text{Receiver}]5\text{h})/[\text{Initial}]0\text{h}$. Compounds were categorized as having high (bound fraction $>90\%$), moderate (50-90%), or low ($<50\%$) plasma protein binding.

Together, these data indicate that TP1 exhibits moderate plasma protein binding in human plasma ($F_u = 11.4\%$), supporting adequate free-drug availability at pharmacologically relevant concentrations, while its plasma protein binding in rat could not be established in this assay, an important caveat when extrapolating rat in vivo pharmacokinetic and pharmacodynamic findings to predicted human free-drug exposure<sup>8-10</sup>.

**Kinetic Aqueous Solubility.** The kinetic aqueous solubility of TP1 was determined in phosphate-buffered saline (PBS, pH 7.4) by a DMSO-dilution shake-flask method with LC-MS/MS quantification. An aliquot (10 μL) of a 10 mM DMSO stock solution of TP1 was added to 990 μL of PBS (pH 7.4) to give a nominal test concentration of 100 μM (final DMSO concentration, 1% v/v). Samples were shaken at 1000 rpm for 1 h at room temperature to allow equilibration and precipitation of any undissolved compound. Samples were then centrifuged (12,000 rpm, 10 min) to pellet undissolved particulate, and the supernatant was collected for analysis. An 8-point calibration curve (0.5-150 μM) was prepared for each compound in 70% acetonitrile from a matched DMSO/organic stock solution. Both calibration standards and test samples were diluted with acetonitrile:methanol (1:1, containing internal standard) prior to LC-MS/MS analysis. Kinetic solubility was calculated by interpolation against the calibration curve and corrected for the 1:1 dilution factor applied during sample preparation. All incubations were performed in triplicate (n = 3)<sup>11-13</sup>.

**Caco-2 Cell Permeability.** Bidirectional permeability of TP1 was assessed across Caco-2 cell monolayers, a standard in vitro model for predicting human intestinal absorption, with atenolol (low permeability marker), propranolol (high permeability marker), and digoxin (P-glycoprotein [P-gp] substrate marker) included as reference compounds. Caco-2 cells (passage 40) were seeded onto 96-well transwell plate inserts at $1 \times 10^5$ cells/cm<sup>2</sup> and cultured for 14-28 days to allow formation of a confluent, polarized cell monolayer, with medium changes every 3-4 days. Monolayer integrity was confirmed by transepithelial electrical resistance (TEER) measurement prior to the transport assay.

Test and reference compounds were diluted from 10 mM DMSO stock solutions to a working concentration of 10 μM in transport buffer (Hanks' Balanced Salt Solution [HBSS] with bovine serum albumin) and applied to either the apical (donor, for apical-to-basolateral [A→B] transport) or basolateral (donor, for basolateral-to-apical [B→A] transport) chamber. Permeation was assessed over

120 min at 37°C, 5% CO<sub>2</sub>, and 95% relative humidity, with all conditions performed in duplicate. Compound concentrations in donor and receiver chambers were quantified by LC-MS/MS.

The apparent permeability coefficient (P<sub>app</sub>) was calculated as: $P_{app} \text{ (cm/s)} = (dC_r/dt) \times V_r / (A \times C_0)$, where dC<sub>r</sub>/dt is the rate of change in receiver chamber concentration, V<sub>r</sub> is the receiver chamber volume (0.1 mL apical, 0.3 mL basolateral), A is the monolayer surface area (0.143 cm<sup>2</sup>), and C<sub>0</sub> is the initial donor chamber concentration. The efflux ratio (ER) was calculated as P<sub>app</sub>(B→A)/P<sub>app</sub>(A→B). Percent recovery was calculated from donor and receiver chamber concentrations at the end of the incubation. Compounds were classified as having low (P<sub>app</sub> A→B < 2 × 10<sup>-6</sup> cm/s), moderate (2-10 × 10<sup>-6</sup> cm/s), or high (> 10 × 10<sup>-6</sup> cm/s) permeability. A compound was classified as a potential P-gp substrate if its efflux ratio without a P-gp inhibitor was greater than 2, and this ratio was reduced by more than 50% in the presence of a P-gp inhibitor<sup>14-16</sup>.
